# Leveraging chromatin accessibility for transcriptional regulatory network inference in T Helper 17 Cells

**DOI:** 10.1101/292987

**Authors:** Emily R. Miraldi, Maria Pokrovskii, Aaron Watters, Dayanne M. Castro, Nicholas De Veaux, Jason A. Hall, June-Yong Lee, Maria Ciofani, Aviv Madar, Nick Carriero, Dan R. Littman, Richard Bonneau

## Abstract

Transcriptional regulatory networks (TRNs) provide insight into cellular behavior by describing interactions between transcription factors (TFs) and their gene targets. The Assay for Transposase Accessible Chromatin (ATAC)-seq, coupled with transcription-factor motif analysis, provides indirect evidence of chromatin binding for hundreds of TFs genome-wide. Here, we propose methods for TRN inference in a mammalian setting, using ATAC-seq data to influence gene expression modeling. We rigorously test our methods in the context of T Helper Cell Type 17 (Th17) differentiation, generating new ATAC-seq data to complement existing Th17 genomic resources (plentiful gene expression data, TF knock-outs and ChIP-seq experiments). In this resource-rich mammalian setting, our extensive benchmarking provides quantitative, genome-scale evaluation of TRN inference combining ATAC-seq and RNA-seq data. We refine and extend our previous Th17 TRN, using our new TRN inference methods to integrate all Th17 data (gene expression, ATAC-seq, TF KO, ChIP-seq). We highlight newly discovered roles for individual TFs and groups of TFs (“TF-TF modules”) in Th17 gene regulation. Given the popularity of ATAC-seq, which provides high-resolution with low sample input requirements, we anticipate that application of our methods will improve TRN inference in new mammalian systems, especially *in vivo*, for cells directly from humans and animal models.

## Introduction

Advances in genome-scale measurement and mathematical modeling herald opportunities for high-quality reconstruction of transcriptional regulatory networks (TRNs). TRNs describe the control of gene expression patterns by transcription factors (TFs) (Hecker et al. 2009; Chai et al. 2014; Marbach et al. 2012), providing mechanistic (and often genome-wide) insight into the complex regulation of cellular behavior (Bonneau et al. 2007). Measurements of chromatin state represent one such advance for TRN inference. For example, chromatin-immunoprecipitation with DNA microarray (ChIP-chip (Ren et al. 2000)) or sequencing (ChIP-seq (Robertson et al. 2007)) enable identification of an individual transcription factor (TF)’s binding sites genome-wide. ChIP experiments provide evidence for TF-target-gene interactions based on proximity of the TF binding site to the gene locus and have proved valuable for TRN inference (Lee et al. 2002; Ouyang et al. 2009; Ciofani et al. 2012). However, their utility is limited in cell types and settings (e.g., physiological) where sample material and/or a priori knowledge of key transcriptional regulators is scarce.

Genome-scale chromatin accessibility measurement (by Formaldehyde-Assisted Isolation of Regulatory Elements (FAIRE)-seq (Giresi et al. 2007), DNase1 hypersensitivity (DHS) (e.g., DNase1-seq) (Xi et al. 2007; Boyle et al. 2008), or most recently Assay for Transposase Accessible Chromatin (ATAC)-seq (Buenrostro et al. 2013)) highlight regulatory regions of the genome that are accessible for TF binding. Similarly, ChIP-seq for histone marks (Barski et al. 2007) (e.g., H3K27Ac, H3Kme3, etc.) correlate with promoters, enhancers, and/or other locus control regions, bound by TFs. Chromatin accessibility and histone-mark ChIP-seq measurements can partially overcome limitations in a priori knowledge of cell-type-specific TF regulators, if integrated with TF DNA-binding motifs (Pique-Regi et al. 2011). Large-scale efforts to characterize individual TF motifs are ongoing, with motifs available for ~700 TFs in human or mouse (~40% coverage for known TFs) (Weirauch et al. 2014; Jolma et al. 2010). Thus, chromatin state experiments integrated with TF analysis provide indirect DNA-binding evidence for hundreds of TFs, the scale of which would be impractical to attain with individual TF ChIP-seq experiments. Of techniques available, ATAC-seq best overcomes limitations in sample material availability, requiring two orders of magnitude fewer cells than a typical ChIP-seq, FAIRE-seq, or DHS experiment in standard, widely-adopted protocols. ATAC-seq is also possible at the resolution of a single cell (Buenrostro et al. 2015).

In the context of TRN inference, chromatin state measurements provide an initial set of putative TF-gene interactions, based on evidence of TF binding near a gene locus; evidence can be direct (TF ChIP-seq) or indirect (e.g., TF motif occurrence in accessible chromatin) and can be used to refine gene expression modeling (Blatti et al. 2015; Wilkins et al. 2016; Qin et al. 2014). Integration of chromatin state data in TRN inference could mitigate false-positive and false-negative TF-gene interactions expected from chromatin-state data analyzed in isolation (Äijö and Bonneau 2016; Siahpirani and Roy 2016). Sources of false positives and negatives include the following: (1) non-functional binding, (2) long-range interactions between genes and regulatory regions (3) the limited availability of individual TF ChIP experiments and incomplete knowledge of TF DNA-binding motifs, and (4) non-bound accessible motifs. Thus, an initial TRN derived solely from chromatin state data can be considered a useful but noisy prior, to be integrated with other data types for TRN inference.

Genome-scale inference of TRNs in mammalian settings is an outstanding challenge, given the increased complexity of transcriptional regulatory mechanisms relative to simpler eukaryotes. Thus, chromatin state measurements are especially important for mammalian TRN inference. Construction of a genome-scale TRN for *in vitro* T Helper Cell Type 17 (Th17) differentiation provided a proof-of-concept for this idea (Ciofani et al. 2012). Rich genomics datasets informed the Th17 TRN: 156 RNA-seq experiments (including knock out (KO) of 20 TFs), ChIP-seq of 9 key Th17 TFs, and microarray from the Immunological Genome Project (Heng et al. 2008). We used the *Inferelator* algorithm (Bonneau et al. 2006; Madar et al. 2010) to infer TRNs from the RNA-seq and microarray data, and independent methods to build networks from TF ChIP and KO data. We showed that rank combination of the networks performed best recovering known Th17 genes and GWAS disease genes associated with Th17 pathologies.

Since our previous publication of the Th17 TRN, the *Inferelator* algorithm underwent several key developments that improve inference in unicellular organisms and are expected to improve TRN inference in a mammalian setting (Greenfield et al. 2013; Arrieta-Ortiz et al. 2015). While the *Inferelator*’s core model of transcriptional regulation still describes differential gene expression as a sparse multivariate linear function of TF activities, the methods used to solve for the TF-gene interaction terms as well as techniques for TF activity estimation have changed. For example, the current version of the *Inferelator* uses a Bayesian approach to incorporate prior information (G-priors with Bayesian Best Subset Regression (BBSR)) (Greenfield et al. 2013). This framework has several desirable features. For example, a TF-gene edge in the prior will not be incorporated in the final TRN without support from the gene expression data, and new edges (not included in the prior) can be learned if there is sufficient support. The current “*Inferelator*-BBSR” presents a promising means to leverage a noisy, chromatin-state-derived prior network in a mammalian setting.

The focus of this work is development of mammalian TRN inference methods from chromatin accessibility and gene expression, data types available or likely feasible for an ever-growing number of cell types and biological conditions. In the context of mammalian TRN inference, several studies build TRNs directly from chromatin accessibility without further refinement by multivariate gene expression modeling (Neph et al. 2012; Rendeiro et al. 2016). Several other studies leverage variance in paired RNA-seq and ATAC-seq datasets; these TRN methods are exciting developments but require that ATAC-seq data be available for all or most RNA-seq conditions (Karwacz et al. 2017; Ramirez et al. 2017; Duren et al. 2017). In contrast, the present work is geared for TRN inference from RNA-seq and ATAC-seq, where ATAC-seq need not exist for more than one gene expression condition.

Development of any TRN inference method requires a comprehensive benchmark with a realistic experimental design, a recurrent challenge in computational biology. To address this challenge and enable rigorous benchmarking of our methods, we extended the genomics resources previously developed by our labs for Th17 TRN inference, using the same Th17 differentiation protocol to generate new ATAC-seq samples. In addition, we augment our initial set of 153 RNA-seq experiments with additional (including unpublished) RNA-seq experiments for a total of 254 *in vitro* Th17 and other T Helper (Th) Cell RNA-seq experiments. Using the RNA-seq and ATAC-seq as input to TRN methods, the quality of resulting TRNs is quantified with precision-recall of “gold-standard” TF-gene interactions in Th17 cells (supported by TF KO and ChIP data (Ciofani et al. 2012; Yosef et al. 2013)). As an alternative to performance relative to a gold standard (unavailable for most cell types), we test TRN quality with out-of-sample gene expression prediction. After extensive benchmarking and method development, we infer an updated TRN model for Th17 cells from all available data (RNA-seq, ATAC-seq, ChIP-seq and TF KO network). Thus, in addition to methods and a genomics resource for mammalian TRN construction, we provide new predictions about transcriptional regulation in Th17 cells, easily accessible to the community in an interactive Jupyter notebook framework.

## Results

### Construction of Th17 benchmark for TRN inference from ATAC-seq and RNA-seq

To test the feasibility of TRN inference from chromatin accessibility and gene expression alone, we generated an ATAC-seq dataset in Th17 cells and other *in vitro* polarized T cells, matching a subset of experimental conditions and time points from the original publication (Ciofani et al. 2012) (Fig. 1A). The ATAC-seq samples displayed characteristic nucleosome-length periodicity in fragment distribution and TSS-localized signal (**Supplemental Fig. S1**). We identified 63,049 accessible regions (peaks), and clustering revealed that most dynamically changed over the T-Helper-polarization time courses (from naïve CD4 T-cell to Th0, Th17, Th2 or T_reg_) (**Supplemental Fig. S2**). These patterns are also apparent from Principal Component Analysis (PCA) (Fig. 1A). Time was the most important driver of chromatin accessibility patterns; the first principal component (PC) explained 55% of the variance and captured peaks changing from two to 48hrs in Th17, Th0, Th2, T_reg_. The second PC captured accessibility differences between Th17 and the other T-cell polarization programs. The Th17 ATAC-seq dataset also contains several perturbation conditions, including TF KO of *Stat3* and *Maf* for Th17 and Th0 conditions (48hr). STAT3 is required for Th17 differentiation and *Stat3* KO dramatically altered Th17 chromatin accessibility, leading to a Th0-like profile (Fig. S2, 1A, red arrow), while *Maf* KO clustered with Th17 (Fig. S2, 1A, gray arrow). Addition of other Th17-polarizing cytokines (IL-1β, IL-23, and/or IL-21) to the Th17 differentiation media (TGF-β and IL-6) did not dramatically change chromatin accessibility patterns at 48 hours.

To the 153 RNA-seq experiments from the original publication, we added an additional 101 RNA-seq experiments, both published (13) and unpublished (88), for a total of 254 experiments (**Fig. 1B, Methods**). The majority (166 samples) were Th17, spanning 1hr to 96hrs, involving KO, siRNA knock down, and/or drug inhibitors of TFs and signaling molecules. The study design also included other T cell polarizations (Th0 (53), Tr1 (9), and Th1/Th2/Treg (2 each)) as well as naïve CD4 T cells (25). Mirroring chromatin accessibility patterns, PCA of the gene expression data revealed time and T-Cell polarization conditions to be important drivers of transcriptional variation (**Fig. 1B, Supplemental Fig. S3**).

While gene expression data are the only required input for the *Inferelator* algorithm (**Methods**), we hypothesized that the inclusion of ATAC-seq data could improve TRN inference. As described above, integration of ATAC-seq with TF motifs can provide indirect evidence for TF-binding events associated with altered chromatin state (**Fig. 1A**) and transcription (**Fig. 1B**). In this study, we generated “prior” networks of TF-gene interactions by associating TFs with putative target genes based on TF motif occurrences within accessible regions near genes (e.g., +/-10 kb gene body, see **Methods). Supplemental Table S1A** provides high-level statistics for two ATAC-seq-derived prior networks: the A(Th17) prior, limited to motif analysis of peaks accessible in 48h Th17 conditions only, and the A(Th) prior, which includes accessible regions from all samples in Fig. 1A. The A(Th17) and A(Th) priors contained ~1.1 and ~2 million putative TF-gene interactions for ~800 TFs. These noisy priors help guide network structure for TRN inference (**Fig. 1C**). Although we do not have an estimate of false edges in our ATAC-seq priors, the study design (**Fig. 1C**) enables quantitative performance evaluation of the resulting TRNs. Specifically, the TRNs are evaluated based on precision and recall of TF-gene interactions from an independent gold standard (G.S.), composed of edges supported by TF KO and/or TF ChIP data. Supplemental Table S1A provides statistics on the G.S. networks and their overlap with other prior information sources. As precision-recall is limited to the TFs previously selected for KO (25 TFs) and/or ChIP (9 TFs) analysis, we also quantify TRN model performance based on out-of-sample gene expression prediction.

**Figure 1.**
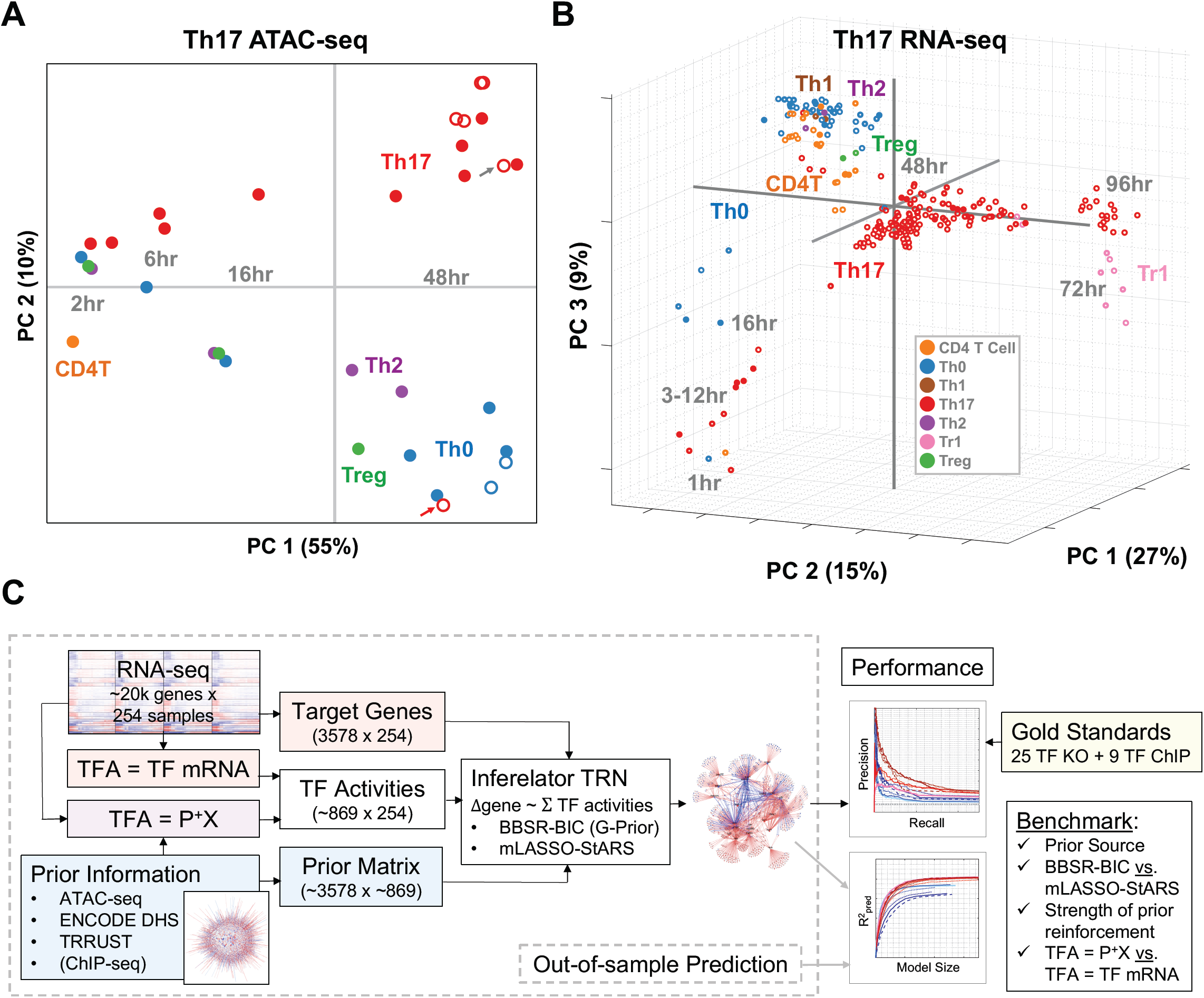
(A) PCA of Chromatin Accessibility Profiles. The 33 ATAC-seq samples are plotted as a function of robustly normalized ATAC-seq peak intensities in PCA space, using the reference set of 63,049 ATAC-seq peaks identified. Open circles denote experimental conditions that deviate from the standard T Cell differentiation conditions (e.g., genetic deletion, additional cytokines). Gray and red arrows indicate Maf and Stat3 KO Th17 conditions, respectively. **(B) PCA of Gene Expression Profiles.** The 254 RNA-seq samples are plotted as a function of all genes in PCA space. **(C) Study Design.** The gene expression data is used to construct both a target gene matrix and to estimate TFA, either with the prior (TFA=P+X) or independently of the prior (TFA=TF mRNA). The *Inferelator* takes target gene and TFA matrices as input and builds models gene expression as a function of TFAs, using either BBSR-BIC or mLASSO-StARS methods. The prior matrix is an optional *Inferelator* input that can be used to reinforce TF-gene interactions with support from the prior. The quality of resulting TRNs is measured using (1) precision-recall of edges in the gold standard and (2) out-of-sample prediction error. The design enables quantitative comparison of prior information source, TFA estimation technique, *Inferelator* method, and strength of prior reinforcement.

Using precision-recall and out-of-sample prediction metrics, we evaluate the effects of several key modeling decisions. Fig. 1C outlines inputs to the *Inferelator* algorithm. From the gene expression dataset, we seek to model the expression patterns of 3578 “target” genes (**Methods**) as functions of TF activities (TFAs). Protein TFAs are rarely measured and technically infeasible for most TRN experimental designs. Thus, the TF activity can be considered a hidden (or latent) variable in the context of TRN inference (Liao et al. 2003; Fu et al. 2011). The most common estimate of TFA is TF mRNA. However, many TF transcriptional activities require protein post-translational modification, and, in those cases, TF mRNA can be a poor proxy for protein transcriptional activity. Thus, TFA estimation based on prior knowledge of TF target genes provides an alluring alternative, as it appears to be technically feasible – requiring only partial a priori knowledge of TF-gene interactions and gene expression data (**Methods**). TFA estimation based on target gene expression quantitatively improved TRN inference using highly curated, literature-based priors in unicellular organisms (Arrieta-Ortiz et al. 2015; Tchourine et al. 2018). However, “prior-based” TFA estimation (from ATAC-seq and other data sources) has yet to be quantitatively evaluated in a mammalian setting at scale.

We test two methods for model selection (i.e., TF-gene interaction terms to include in the TRN models): (1) the most recently published version of *Inferelator* (Bayesian Best Subset Regression with Bayesian Information Criteria for model selection (BBSR-BIC) (Arrieta-Ortiz et al. 2015)) and (2) an alternative proposed here, modified Least Absolute Shrinkage and Selection Operator (Studham et al. 2014; Gustafsson et al. 2015) with Stability Approach to Regularization Selection (Liu et al. 2010) (mLASSO-StARS), to overcome challenges inherent to mammalian TRN inference. Specifically, we hypothesized that mLASSO-StARS would (1) scale better with the increased transcriptional complexity expected in a mammalian setting (e.g., larger models) and (2) stability-based StARS model selection would perform better for relatively smaller mammalian gene expression datasets (10s-100s of samples versus 1000s of samples in the unicellular organisms for which *Inferelator* BBSR-BIC was originally developed) (**Methods**). Thus, we compare mLASSO-StARS and state-of-the-art BBSR-BIC for mammalian TRN inference (Arrieta-Ortiz et al. 2015). Notably, StARS model selection has been used in several related biological contexts already (Kurtz et al. 2015; Zhang et al. 2016; Caballe Mestres et al. 2017).

Prior information can enter our inference procedure at two steps (1) prior-based TFA estimation (described above) and (2) to reinforce prior-supported TF-gene interactions (edges) at the multivariate regression step, using either BBSR-BIC or mLASSO-StARS (**Fig. 1C, Methods**). The strength of prior reinforcement controls the relative contribution of prior evidence (e.g., TF ChIP, ATAC-seq motif analysis) to evidence from the gene expression model (variance explained by individual TFs). Effectively, the strength of prior reinforcement determines the frequency with which prior edges enter the network relative to edges supported by gene expression modeling alone. The appropriate level of prior reinforcement depends on (1) the quality of the prior information and (2) quality and number of gene expression samples available for model building. Clearly, the strength of prior reinforcement will influence TRN inference; thus, we test several levels of reinforcement in our study design. We also test several sources of prior information, in addition to ATAC-seq.

### Modified LASSO-StARS improves inference of a mammalian TRN

As illustrated in **Fig. 1C**, we use precision and recall over multiple benchmark networks to quantitatively evaluate the impact of key modeling decisions on resulting Th17 TRN models (prior sources, method of TFA estimation, model selection method (BBSR-BIC or mLASSO-StARS), and strength of prior reinforcement). Both BBSR-BIC and mLASSO-StARS provide confidence estimates for every TF-gene interaction (“edge”) predicted (**Methods**), and these confidences can be used to prioritize TF-gene interactions and rank edges. For each confidence cutoff, predictions can be evaluated in terms of precision (fraction of predicted edges in the gold standard) and recall (fraction of the total gold standard recovered). There is a well-known trade-off between precision and recall. In general, a high confidence cutoff (yielding a small network) favors high precision but low recall, while a low confidence cutoff (many interactions included in the network) favors high recall at the cost of lower precision. Therefore, it is instructive to plot precision and recall as a function of confidence cutoffs, as in **Fig. 2A**. Calculating the area under the precision-recall curve (AUPR) provides a useful summary of the precision-recall curve.

**Figure 2.**
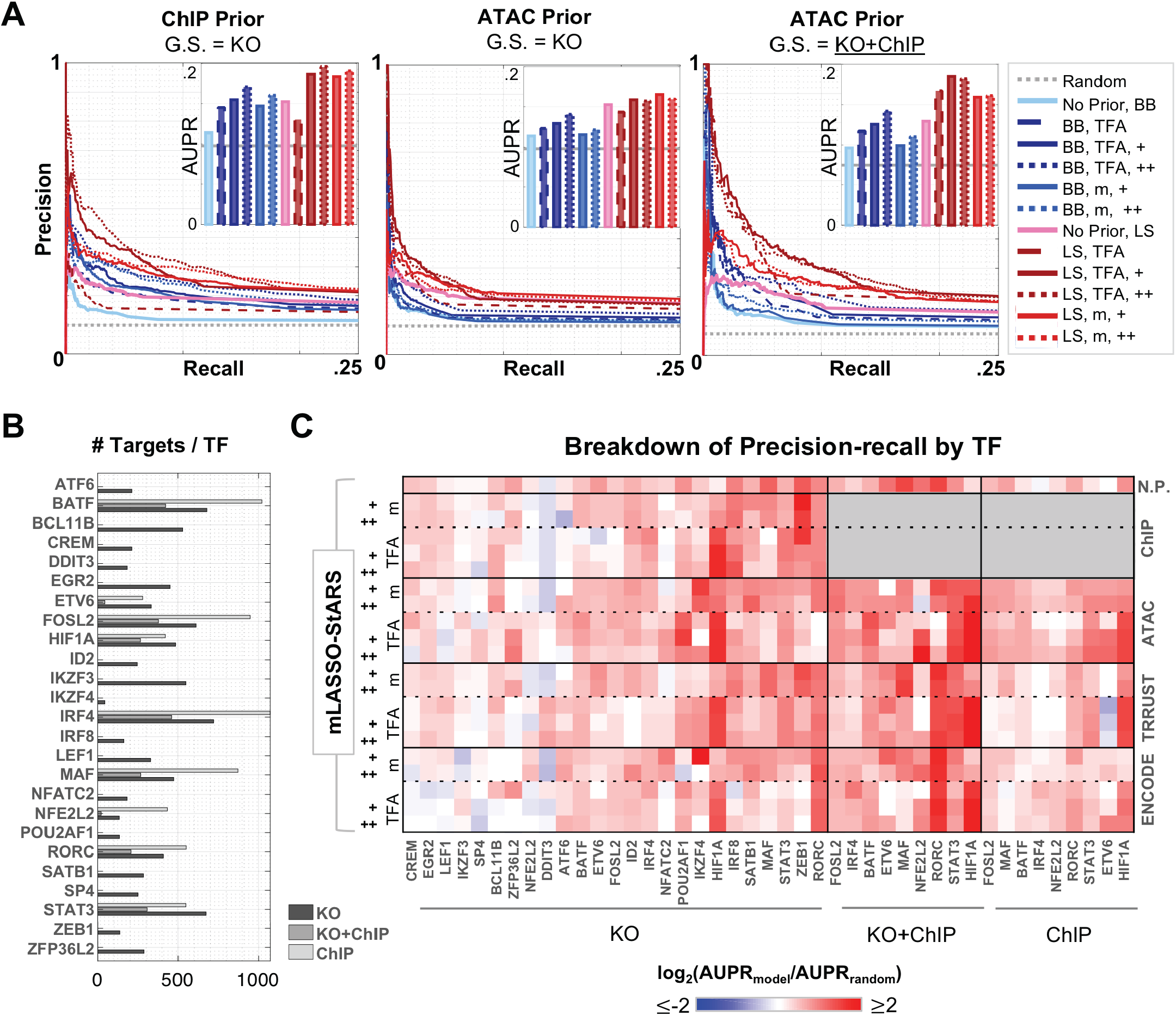
(A) Precision-recall of Th17 TRNs. The left two panels enable comparison of TRNs built from ChIP versus Th17 ATAC priors, quantified by precision-recall of the KO gold standard (G.S.) (25 TFs, 8875 interactions), insets display AUPR. The performance of several TRNs are plotted for each prior, based on Inferelator method (LS = mLASSO-StARS (reds), BB = BBSR-BIC (blues)), TFA estimate (m = TF mRNA, TFA = P^+^X), and “+” corresponds to strength of prior reinforcement. Random and “No Prior” control TRNs serve as references in both panels. The right panel shows precision-recall of the KO-ChIP G.S. (9 TFs, 2375 edges) for TRNs built from the ATAC prior. **(B) Number of targets per TF in the Gold Standards.** Targets per TF are limited to the 3578 considered by the model. **(C) TF-specific TRN performance.** For each G.S., AUPRs were calculated for each TF individually. TF-specific performance of TRNs is quantified as the log_2_-foldchange between AUPR of the TRN model relative to random. “+” indicates strength of prior reinforcement, and “m” and “TFA” denote TFA estimation methods as in **(A)**.

The quality of precision-recall analysis is dependent on the quality of the gold standard (G.S.). In this study, we have access to 25 TF KO RNA-seq experiments (Th17 48h time point), from which TF-target gene interactions were derived from differential expression analysis (Yosef et al. 2013; Ciofani et al. 2012). There are also 9 TF ChIP-seq experiments (Th17 48h time point), from which TF-gene interactions were derived based on ChIP-peak proximity to gene locus (Ciofani et al. 2012). However, gold standards derived from either datatype have caveats. For example, differential analysis of TF KO expression profiles is imperfect, as cellular TRNs adapt to the original TF perturbation over time, leading to partial compensation by paralogs (false negatives) and changes in expression of TFs and signaling molecules downstream of the original TF (false positives). The TF ChIP G.S. will also contain false positives (TF ChIP peaks are not necessarily functional) and false negatives (without 3D-chromatin conformation in Th17 cells, we limit to linear distances between TF ChIP peak and gene loci). Generating a gold standard from edges supported by both TF KO and TF ChIP reduces false-positive rate, but it does so at the expense of false negatives.

For each G.S. information source, **Supplemental Table S1A** summarizes the number of edges, TFs, and target genes as well as percent overlap with other prior information sources. **Fig. 2B** shows the TF degree (number of gene targets) in each G.S. Because we have KO data for 25 TFs but KO+ChIP data for only 9 TFs, we also include precision-recall analysis of the KO G.S. As expected, TF ChIP data improves Th17 TRN inference (**Fig. 2A**, left panel). The cases explored include prior-based TFA estimation and TF-mRNA-based TFA (with moderate and high prior reinforcement). For all but one case (prior-based TFA without reinforcement), mLASSO-StARS methods outperform the “equivalent” BBSR-BIC method. We tested inference with the ATAC priors (**Fig. 2A** central and right panels, **Supplemental Fig. S4**), comparing to the ChIP-seq derived prior on recovery of the KO G.S. The ATAC prior boosts performance relative to the “No Prior” control TRNs, but the effect is smaller, likely reflecting increased levels of noise in the ATAC-seq prior (as described in **Introduction**). mLASSO-StARS methods with the ATAC prior show better precision-recall on the likely higher-quality G.S. (KO+ChIP, **Fig. 2A** right panel). In contrast to results with a ChIP prior, increasing the strength of prior reinforcement from moderate to high yields no advantage for the noisier ATAC prior. This suggests ATAC-seq prior reinforcement should be limited to moderate rather than high (i.e., gene expression data should be relied on to select a small subset of the regulatory hypothesis from the ATAC-seq prior network). For similar levels of prior reinforcement, prior-based TFA models outperform TF mRNA at low recall, and this trend holds for the other priors tested below. For all ATAC TRNs and both gold standards, mLASSO-StARS outperforms BBSR-BIC.

To explore experimental designs without context-specific chromatin accessibility, we tested integration of two contrasting, publicly available prior information sources. The first is TF motif analysis of ENCODE DHS data from 25 mouse tissues, none of which include Th17 (Stergachis et al. 2014). The second is derived from the curated TRRUST database of human TF-gene interactions (Han et al. 2015). **Supplemental Table S1A** provides high-level statistics on the resulting priors. While the ENCODE DHS prior includes ~1.5 million interactions between 546 TFs and ~17k genes (similar scale to the ATAC priors), the TRRUST prior is sparse: ~7k interactions between 582 TFs and ~2k genes. The TRRUST and ENCODE priors overlap less with the gold standards than context-specific priors, and this is reflected in lower precision-recall, relative to ChIP and ATAC priors (**Supplemental Fig. S4**). However, use of either the ENCODE or TRRUST prior improves performance relative to the No Prior control; this improvement is substantial for prior-based TFA models. Thus, across priors, TFA methods, and levels of prior reinforcement, mLASSO-StARS outperforms BBSR-BIC.

To evaluate performance on experimental designs with fewer gene expression samples, we reduced the gene expression matrix from 254 to 50 randomly selected samples (**Supplemental Fig. S6**). The reduced sample size had minor impact on precision-recall, especially for context-specific ChIP and ATAC priors, suggesting that these priors significantly guided TRN learning. Most conditions (60%) have one or more biological replicates, and this might also contribute to the hardly-diminished performance using smaller gene expression datasets. These results bode well for extension of our methods to cellular contexts where gene expression data are less abundant.

We also evaluated methods at the resolution of individual TFs, to evaluate how the different modeling decisions affect target prediction for each TF (**Fig. 2C**). There is nearly an order-of-magnitude difference in degree per TF in the KO+ChIP G.S. (**Fig. 2B**), so this per-TF analysis additionally ensured that results were not dominated by a few high-degree TFs. Overall, mLASSO-StARS also outperformed BBSR-BIC at TF-specific AUPR resolution (**Supplemental Fig. S7**).

AUPRs for many TFs were dependent on TFA estimation procedure (**Fig. 2C**). TF mRNA can be a good estimate of protein TFA for TFs whose main source of regulation is transcriptional, while, for ubiquitously expressed TFs regulated by posttranslational modification, prior-based TFA would be expected to outperform TF mRNA estimates. Consistent with this, models with prior-based TFA have higher AUPR for STAT3 (ChIP prior with KO G.S., ATAC prior with KO+ChIP and ChIP G.S.), while prior-based TFA did not improve prediction of RORC targets. With the ChIP prior (which includes RORγt TF ChIP), prior-based TFA AUPR was on par with TF mRNA AUPR. However, for the noisier ATAC-seq prior, prior-based TFA performed little better than random, while TF mRNA models (including the “No Prior” control) performed well (all G.S.’s). For ATAC-seq-based TRN inference, target prediction for some TFs (HIF1A, STAT3, NFE2L2) is better using prior-based TFA, while TF mRNA is better for some TFs (RORC, MAF, FOSL2) and roughly equivalent for others. Summarizing across priors and parameter sets, no TFA method dominates (**Supplemental Fig. S7B**). Based on these results, we later generate “final” Th17 TRNs using a combination of prior-based and TF-mRNA TFA estimates.

### Th17 TRN models predict out-of-sample gene expression patterns

We next evaluated whether the TRN models could predict out-of-sample gene expression patterns. In contrast to precision-recall, gene expression prediction does not require a gold standard and provides the opportunity to evaluate all interactions in the model. Thus, gene expression prediction is especially important in poorly characterized cellular systems, for which gold standards do not exist and precision-recall is not possible. We chose three out-of-sample prediction challenges for the TRN models with distinct patterns of gene expression, highlighted in the PCA plot (**Fig. 3A**). These three leave-out sets are: “Early Th17” (all Th17 time points between 1-16 hours, 8 samples), “All Th0” (Th0 samples for all time points and perturbations, 53 samples), or “Late Th17” (18 Th17 samples from 60-108h post-TCR stimulation). For both BBSR-BIC and mLASSO-StARS methods, we built models and tested prediction over a range of edge confidence values and corresponding model sizes (**Methods**). We quantified predictive performance using r-squared of prediction, 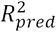whose values range from one (perfect prediction) to -∞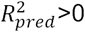 indicates that the model has predictive benefit. We compared predictive performance versus model size for TRNs built with an ATAC-seq prior as well as control (“No Prior”) networks for each leave-out set (**Fig. 3B, 3C**). Across all leave-out sets and methods, out-of-sample prediction improved most as models expanded from average size of 1 TF/gene to 5 TFs/gene (**Fig. 3B**). Most methods performed similarly well from 0-10 TFs/gene, with the exception of BBSR-BIC models using prior-based TFA, where prediction was worse. Predictive performance plateaued at ~10-15 TFs/gene, depending on the leave-out set (Fig. 3B, S8). For model sizes 10-15 TFs/genes, the mLASSO-StARS models outperformed BBSR-BIC models (Fig. 3B, 3C). These results, together with the precision-recall, support mLASSO-StARS methods over BBSR-BIC for mammalian TRN inference. In addition, moderately reinforced mLASSO-StARS models predicted slightly better than the other mLASSO-StARS models in terms of precision-recall and out-of-sample prediction; thus, we use mLASSO-StARS with moderate prior reinforcement to construct “final” Th17 TRNs.

**Figure 3.**
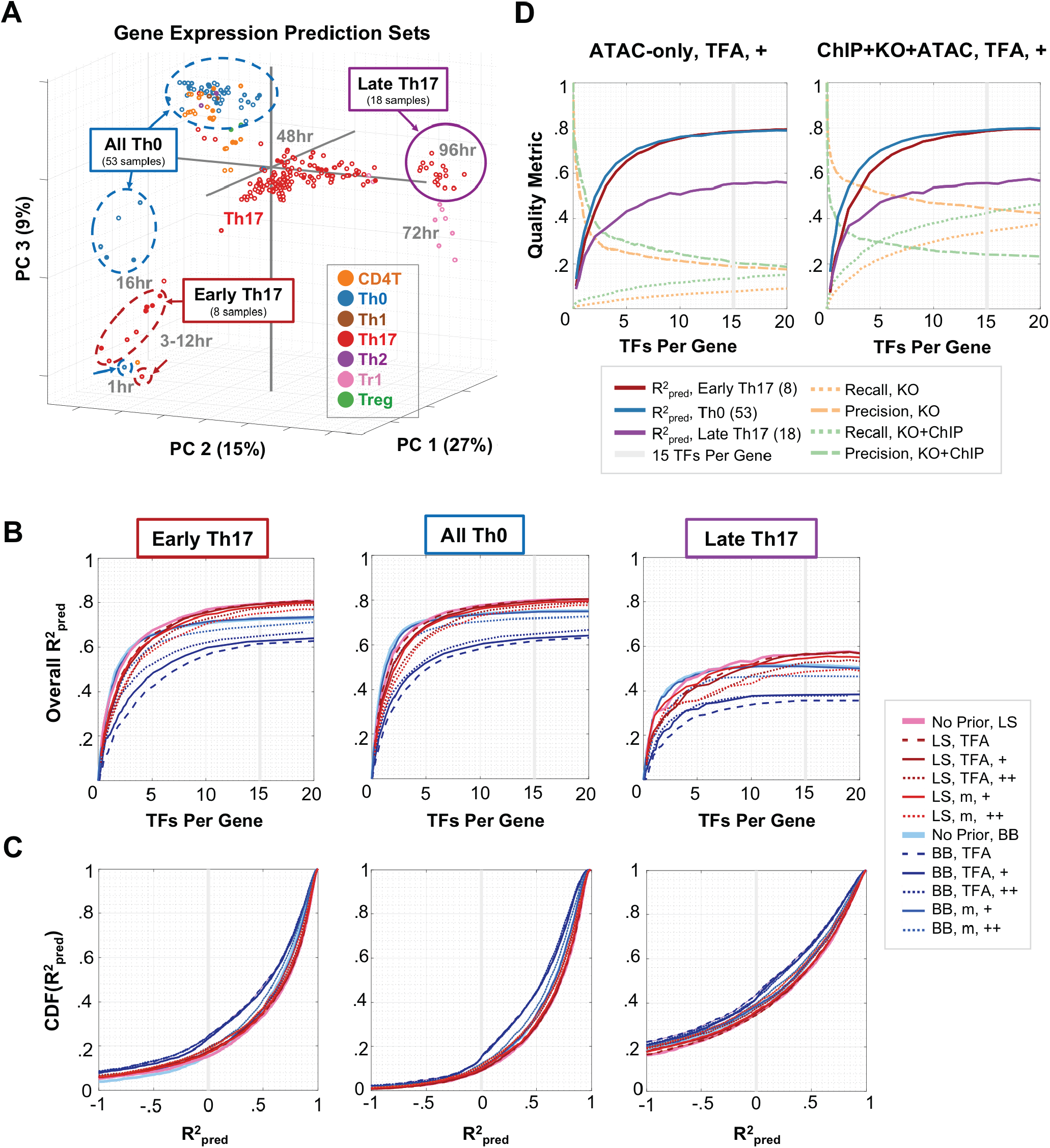
(A) Leave-out sets for gene-expression prediction. Model selection and parameter estimation of TF-gene interactions were performed in the absence of specified leave-out sets (All Th0, Late Th17, and Early Th17), circled in PCA space. **(B) Gene expression prediction by method.** R^2^_pred_ for each leave-out set is plotted as a function of average number of TFs per gene. In the legend, LS = mLASSO-StARS, BB = BBSR-BIC, m = TF mRNA, TFA = prior-based TFA estimate, “+” corresponds to strength of prior reinforcement. Gray line corresponds to a model cutoff of mean 15 TFs per gene. **(C) Distributions of R^2^_pred_ values.** For model-size cutoff of mean 15 TFs/gene, empirical cumulative distribution functions (CDFs) of per-gene R^2^_pred_ values are plotted for each method. **(D) Model quality metrics versus model size.** For two TRN models built with Th17 ATAC (left panel) or ChIP+KO+ATAC (right panel) priors (mLASSO-StARS, bias = .5, TFA = P^+^X), the quality metrics (R^2^_pred_ for each leave-out set, precision and recall) are plotted as a function of model size. The model size used for subsequent analysis is also highlighted.

While we recommend StARS edge stabilities to rank interactions, we used the quality metrics (precision, recall,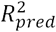) to guide selection of model-size cutoff for the final Th17 TRN; this modeling decision and supporting data are detailed in Methods. These quality metrics are plotted versus model size (**Fig. 3D**) for moderately reinforced prior-based TFA models with the Th17 ATAC (“ATAC-only”) or ChIP+KO+ATAC prior, while results for other conditions can be found in **Supplemental Fig. S9**. The ChIP+KO+ATAC prior (**Methods**) represents our best (combined) source of prior information and will be later used to generate a final Th17 TRN. Once average model sizes reach ~15 TFs/gene (**Supplemental Fig. S8**), predictive performance plateaus, suggesting an average of 15 TFs/gene as a potential cutoff for edge inclusion in the network. Standardizing network sizes to 53k TF-gene interactions (~15 TFs/gene), we calculated the percent edge overlap among TRNs built from ATAC, ChIP, KO, ENCODE DHS or TRRUST priors as well as priors combining ATAC-seq with ChIP and/or KO data (**Methods**). For each prior, we considered five modeling modes: prior-based TFA estimates with no, moderate or strong prior reinforcement and TF mRNA TFA estimates with moderate and strong prior reinforcement. The percentage of shared edges between TRNs ranged from 83% to 10%. We clustered the networks based on overlap to visualize how the modeling decisions affected resulting TRNs on a global scale (**Supplemental Fig. S10**); detailed discussion of these trends is contained in **Supplemental Note S1**.

### “Core” Th17 TRNs contain literature-supported TF-gene interactions

Our primary objective is to assess the feasibility of high-quality TRN inference from gene expression and ATAC-seq data. Therefore, it is important to examine the Th17 TRNs at high resolution. Here, we focus analysis on TRN predictions for 18 “core” Th17 TFs and genes, readily familiar to Th17 biologists (**Fig. 4**). Individual interactions between TFs and genes in this “core” have been the focus of many studies (Christie and Zhu 2014; Li et al. 2014), which we leverage to evaluate the ATAC-based Th17 core TRNs: moderately reinforced TF-mRNA-and prior-based TFA TRNs individually and combined into a final ATAC-only TRN (as described in Methods). To visually compare these networks, we developed jp_gene_viz, a Jupyter-notebook TRN visualization tool. In addition to the ATAC TRNs (**Fig. 4A**), we include the KO-ChIP G.S. (without inference) and the No Prior TRN as references (Fig. 4B, 4C). For more detailed exploration, all 36 LASSO-StARS Th17 TRNs (from **Supplemental Fig. S10**), gold standards, and final, combined TRNs are available in a Jupyter-notebook binder: https://mybinder.org/v2/gh/simonsfoundation/Th17_TRN_Networks/master.

**Figure 4.**
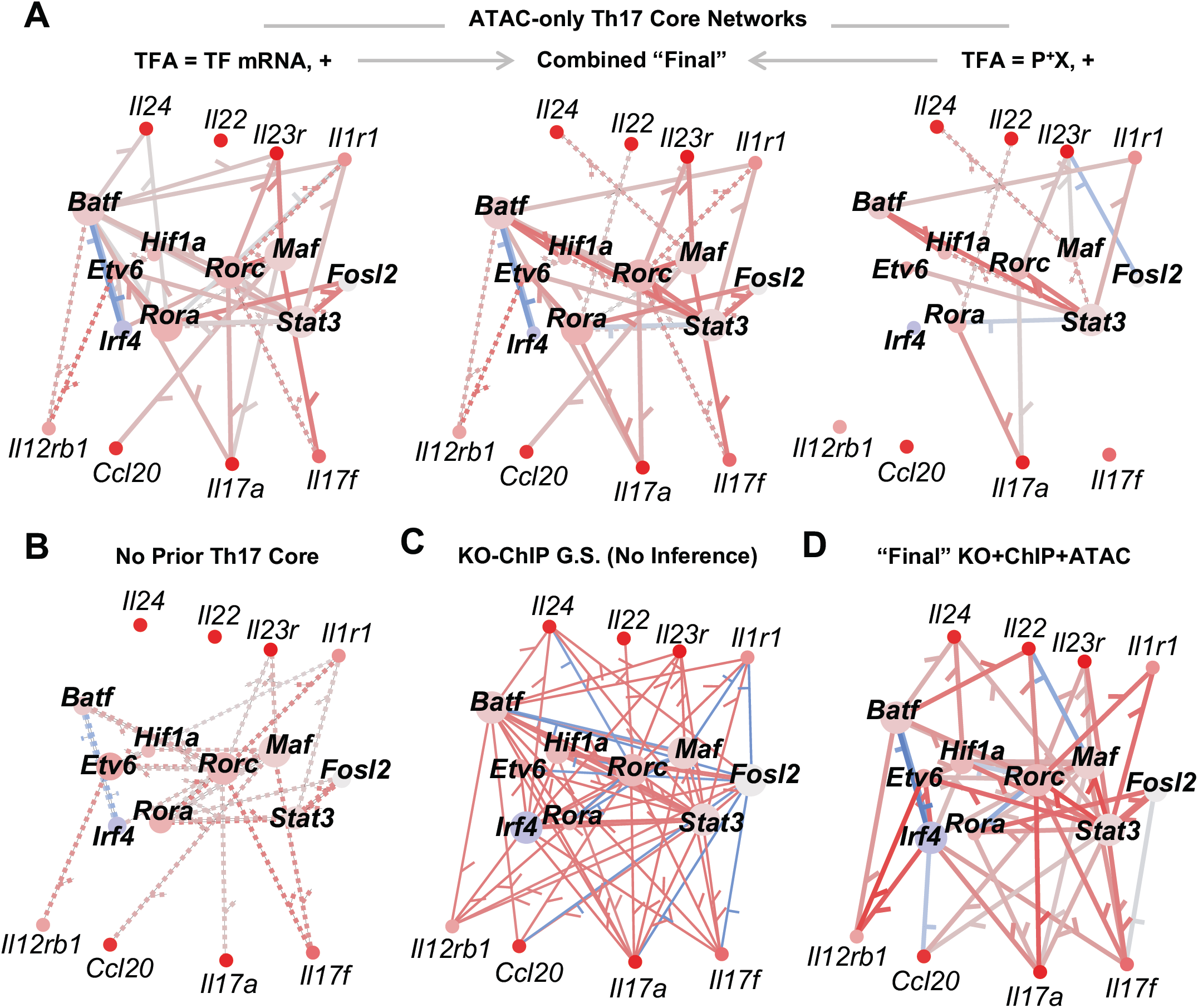
(A-D) Th17 core TRN models. “Core” Th17 genes and TFs were selected from the literature for visual comparison with jp_gene_viz software. Network size was limited to an average of 15 TFs per gene for Inferelator networks using: **(A)** Th17 ATAC prior, **(B)** no prior, and **(D)** ChIP+ATAC+KO prior. The edges in Inferelator TRNs are colored according to partial correlation (red positive, blue negative) and weighted relative to edge stability. **(C)** represents the full KO-ChIP G.S. from Ciofani et al., where edge sign is based on differential gene expression analysis between TF KO and control and edge weight is constant. Nodes are colored according to gene expression at 48h in Th17, z-scored relative to Th17 and other T Helper cell time points (red/blue = increased/decreased expression). The “Final” KO+ChIP+ATAC **(D)** and ATAC-only **(B)** TRNs max-combine networks built using TF mRNA and prior-based TFA (**Methods**).

From the literature and the KO-ChIP G.S. (**Fig. 4C**), there is support for edges between RORC and several key Th17 cytokines and receptors: *Il17a, Il17f, Il22, Il1r1* and *Il23r*. Two of these interactions (*Il17a, Il23r*) were present in the ATAC-seq prior and both TF-mRNA-and prior-based TFA ATAC TRNs (Fig. 4A, left and right). Four of the interactions (*Il17a, Il17f, Il1r1* and *Il23r*) are recovered on the basis of gene expression modeling alone, as they were recovered by the No Prior TRN and also the TF-mRNA ATAC TRN. Interestingly, the prior-based TFA ATAC TRN (right panel Fig. 4A) recovers a data-driven edge between RORC and *Il22*. By combining predictions from both ATAC TRNs (see **Methods**), all five RORC targets are recovered (central panel, Fig. 4A). As expected, inclusion of RORC ChIP and/or KO in the prior leads to recovery of all five TF targets as well (**Fig. 4D**).

STAT3 is required for Th17 differentiation, playing a crucial role in the activation of *Rorc*. There is support for this interaction not only from context-specific ATAC-seq prior but also the ENCODE DHS prior. Consistent with the poor quality of mRNA-based STAT3 predictions (**Fig. 2C**), this interaction is not present in the No Prior TRN, but is recovered in ATAC and ENCODE TRNs, even the TF mRNA models, for which prior reinforcement likely overcomes weak correlation between TF mRNA and protein activity level (Fig. 4A-C, **S11**).

MAF is another key regulator of Th17 cytokine and receptor expression, with KO-ChIP G.S. edge support for *Il17a, Il17f, Il23r*, and *Il1rb*. There is ATAC-seq support for MAF regulation of *Il17a, Il17f, Il23r*. The prior-supported targets are recovered by the TF-mRNA-based ATAC models, but only the *Il23r* interaction is present in prior-based TFA models (**Fig. 4A**). Similar to RORC, this suggests the signal between *Maf* mRNA and target genes might have been better than the ATAC-inferred MAF TFA. Prior reinforcement also played a role, as only two of the four interactions are present in the No Prior TRN (**Fig. 4B**). In the absence of context-specific prior information and a strong signal from the gene expression model, only a single edge (*Il23r*) was recovered by one of the ENCODE models (**Supplemental Fig. S11**).

These results highlight the potential for TRN inference in new settings, where integration of chromatin accessibility and gene expression is more feasible than sequential TF ChIP and KO experiments. Consistent with the TF-resolved AUPR analysis (**Fig. 2C**), they also suggest that there is value to building models from both TFA methods. For construction of the “final” Th17 TRN, we combine models based on both TFA methods, choosing only moderate levels of prior reinforcement to accommodate useful but noisy ATAC-seq data in our final combined prior. The literature-curated core of our final Th17 TRN contains the RORC and MAF cytokine and receptor interactions highlighted from the literature, as well as the established connection between STAT3 and *Rorc* (**Fig. 4D**),

### ATAC-derived Th17 TRNs contain known and novel Th17 transcription factors

Having verified that the Th17 TRNs contain core Th17 TF-gene interactions from the literature, we develop a global, unbiased analysis of the final ChIP+ATAC+KO TRN to identify new network predictions coordinating relevant Th17 biology. In addition, we extend our analysis to a final TRN using the ATAC-only prior, to simulate mammalian TRN inference in less well-studied systems, where KO and/or ChIP data might not be available. Overall, prior-supported edges make up 63% and 43% of the ~53k TF-gene interactions in ChIP+KO+ATAC and ATAC-only TRNs, respectively (**Supplemental Table S5**). Of the 715 potential TF regulators considered for final models, nearly all (~95%) have targets in the final TRNs, with positive interactions outnumbering negative nearly two-fold (1.8:1 (ChIP+KO+ATAC) or 1.9:1 (ATAC-only)). TF degree varies dramatically (**Supplemental Fig. S12, S13**). Whereas the ATAC-only network democratizes TF degree distribution (no TF has > 500 targets), the addition of ChIP and KO leads to very high degree for several TFs in the ChIP+KO+ATAC TRN (>500 targets for IRF4, BATF, MAF, SP4, FOSL2, and STAT3). While there is a bias for TFs in the prior to have higher degree, several TFs without prior support have >100 targets in the final networks (4 TFs for the ChIP+KO+ATAC TRN and 14 TFs for the ATAC-only TRN). Thus, prior information is not weighted so strongly as to preclude inclusion of the many TFs without known motifs. This aspect is important for discovery and also holds for TFs with edges in the prior, too. While only 16% or 7% of input prior edges remain in the TRN, 27% or 46% of learned regulatory interactions for TFs with prior information are new (not originally in the prior) in KO+ChIP+ATAC or ATAC-only TRNs, respectively). Thus, in addition to reducing false positives, our method can also reduce false negatives. For example, motif analysis of the RORC ChIP data revealed that only ~1/3 of RORC peaks contained a RORC motif (at motif occurrence cutoff P_raw_=1E-4). Although RORC can bind DNA directly, the RORC ChIP data suggests that RORC might also bind DNA indirectly (e.g., via TF complexes). Such indirect binding would be difficult to detect by ATAC-seq motif analysis alone, but their gene targets can be recovered via gene expression modeling.

**Figure 5.**
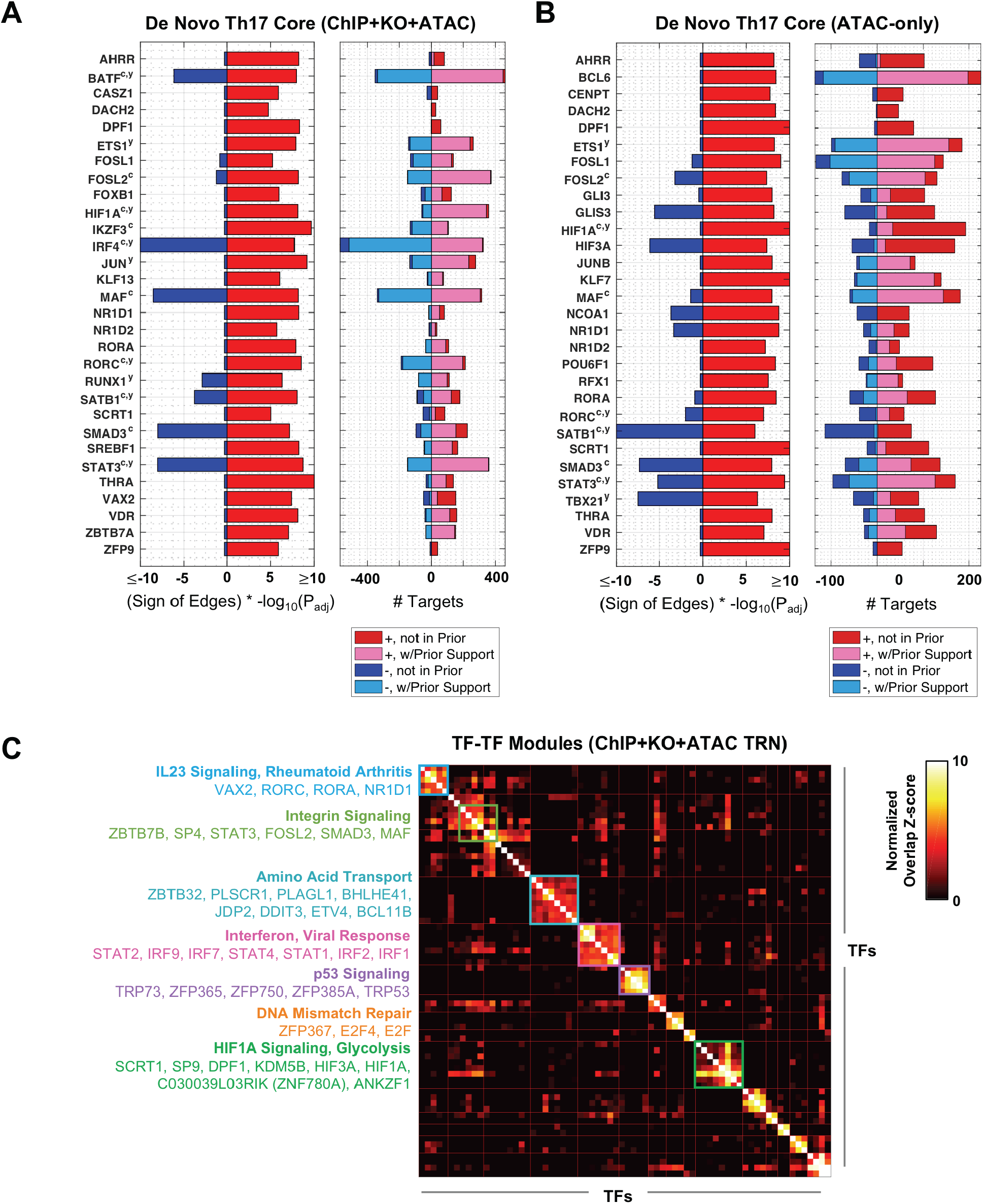
De Novo Th17 Core TFs in the ChIP+KO+ATAC TRN (A) and ATAC-only TRN (B). TFs were included as De Novo Th17 Core TFs if, at a FDR = 10%, (1) positive gene targets were enriched for up-regulated Th17 genes or cytokines and receptors or (2) negative gene targets were enriched for down-regulated Th17 genes or cytokines and receptors. Left panel indicates enrichment in these sets. Right panel denotes number, sign and prior support of TF target edges. Superscripts c and y indicate TF Th17 association from (Ciofani et al. 2012), (Yosef et al., 2013), respectively. **(C) Top 15 TF-TF Modules for ChIP+KO+ATAC TRN**. TFs were clustered into modules on the basis of shared <u>positive</u> target genes between TFs (see Methods). The Top 15 TF-TF modules are displayed. Gene-set enrichment was used to annotate clusters, and TF members are listed. Asterisk indicates TF-TF modules that contain Th17-promoting core TFs.

Many of the TFs with highest degree are shared between the ChIP+KO+ATAC and ATAC-only TRNs (e.g., IRF4, BATF, SP4, RXRA, STAT3,… **Supplemental Fig. S12, S13**). We developed an unbiased approach to identify key regulators of the Th17 program. TFs were included in the set of “core” Th17 regulators if they met one of two criteria: (1) The TF promotes Th17 gene expression through activation of Th17 genes (i.e., the TF’s positive edges are enriched in up-regulated Th17 genes) or (2) The TF promotes Th17 expression through repression of non-Th17 genes (i.e., TF’s negative edges are enriched in down-regulated Th17 genes) (**Methods**). This analysis yielded strikingly similar de novo Th17 core TFs for both ChIP+KO+ATAC and ATAC-only TRNs. Fig. 5A,B highlights the Top-30 core TFs with several recognizable Th17-specific TFs from the literature (RORA, RORC, STAT3, MAF) in both networks. We note that this “core” TF analysis is robust to model-size cutoffs, as analysis of TRNs with average model-size 5 or 10 TFs/gene yields similar results (**Supplemental Fig. S14**). Similarly, top-degree TFs per TRN are robust across model sizes (**Supplemental Fig. S15**).

### TF-TF modules exhibit coordinated control of gene pathways in Th17

To aid in exploring the large Th17 TRNs (~53k TF-gene interactions), we designed a TF-TF module analysis to define clusters of TFs with significant overlap in target genes (see **Methods**, Fig. 5C, **S16, S18**). We then applied a comprehensive gene-set enrichment analysis to predict functional roles for the TF-TF clusters, looking for consensus among pathway enrichments from five databases (Gene Ontology, Pathway Commons, KEGG, WikiPathways and signatures from MSigDB (Consortium 2004; Cerami et al. 2010; Kanehisa and Goto 2000; Pico et al. 2008; Liberzon et al. 2011)) (**Supplemental Fig. S17, S19**). Most TF-TF clusters were conserved between ChIP+KO+ATAC and ATAC-only networks, and, within clusters, TFs shared features. For example, several clusters contained TFs defined in the de novo Th17 cores. RORC was a member of a Th17-promoting TF-TF Module including RORA, NR1D1, VAX2 (highlighted with a light-blue square, Fig. 5C, **S16-S19**); functional annotations for this cluster include “IL23 Signaling” and “Rheumatoid Arthritis”, which are consistent with prior knowledge. Th17-promoting genes HIF1A, HIF3A, DPF1, SP9, and SCRT1 cluster with 5-6 other TFs (green square), and enrichments for this cluster include “hypoxia”, “HIF1A transcription factor network”, and “glycolysis”.

Other clusters contained TFs that promote the expression of genes repressed at 48h in Th17 cells. One such cluster contained Th1 TFs (IRF1, STAT1 and STAT2), many other interferon response factors (IRF2, IRF7, IRF9) as well as STAT2 and STAT4 (hot-pink box, Fig. 5C). As expected, this “interferon cluster” has enrichments for “Response to Interferon Gamma”, “Type 1 Interferon Pathway”, “Response to Virus”. Interestingly, although TF gene expression for this cluster is highest in Th1 relative to other T Cell populations at 48hrs, gene expression is at its highest at the 1hr Th17 time point, suggesting an interferon-like response for Th17 cells very early in the Th17 polarization time course (**Supplemental Fig. S16, S18**). This result is consistent with predictions from another Th17 TRN (Yosef et al. 2013), in which authors also predict roles for IRF1, IRF2, IRF9, STAT1, STAT2 within the first four hours of Th17 polarization. Both findings are consistent with potential plasticity, observed *in vivo*, in Th17 cell programs that are homeostatic or pathogenic, with expression of Th1-like features in the latter.

Gene-set enrichment provides functional predictions for other TF-TF Modules, including: (1) “Amino Acid Transport” (ZBTB32, PLSCR1, PLAGL1, BHLHE41, JDP2, DDIT3, ETV4, BCL11B),(2) “Integrin Signaling” (ZBTB7B, SP4, STAT3, FOSL2, SMAD3, MAF), (3) “DNA Mismatch Repair” (ZFP367, E2F4, E2F), (4) “p53 Signaling” (TRP73, ZFP365, ZFP750, ZFP385A, TRP53), and others (Fig. 5C, **S17, S19**). These predictions provide confirmation of TRN quality, as many modules have predicted function in processes for which individual TFs are already implicated (e.g., HIF1A and HIF3A appear in the HIF1A/hypoxia module, TRP53 and TRP73 appear in the p53-signaling module). In addition, TF-TF modules and functional annotations are largely conserved between KO+ChIP+ATAC and ATAC-only TRNs, the latter prior network being much more economically feasible than the first. More fundamentally, these predictions suggest how altering sets of TFs might influence Th17 pathways and responses.

### New phenotypes are associated with TFs in the Th17 TRN

Th17 cells have been implicated in the pathogenesis of multiple autoimmune diseases (Stadhouders et al. 2017). We previously tested whether genes co-regulated by the “Th17 core” (RORC, STAT3, BATF, IRF4 and MAF) were enriched for gene sets from GWAS of nine autoimmune diseases and three “negative controls” (Alzheimer’s, schizophrenia and Type 2 Diabetes) (Ciofani et al. 2012). Consistent with the known role for Th17 in autoimmune disease, genes from seven of the autoimmune-disease sets were enriched (Crohn’s Disease, Type 1 Diabetes, celiac, rheumatoid arthritis, psoriasis, ulcerative colitis and multiple sclerosis) (Ciofani et al. 2012). The known roles of Th17 cells have since expanded to include pathogenesis of obesity-related diseases (Endo et al. 2017; Harley et al. 2014) and psychiatric disorders (Debnath and Berk 2014; Choi et al. 2016). In parallel, the number of genome-wide association studies have grown exponentially (MacArthur et al. 2016), and, as demonstrated above, our network model has improved in both comprehensiveness and accuracy.

We performed an extensive, unbiased GWAS analysis of the final updated (KO+ChIP+ATAC) Th17 TRN, including any phenotype with five or more associated genes. 991 phenotypes met this criterion. Not only did we expand this analysis in terms of phenotypes considered, we more broadly queried the Th17 TRN, testing for the enrichment of GWAS gene sets in the targets of 605 TFs individually (**Supplemental Table S6**). Despite the large number of TF-phenotype associations tested, eight reached significance (FDR = 10%, Fig. 6A). STAT3 targets were significantly enriched for genes associated with inflammatory bowel disease (IBD) as well as the two IBD-subtypes, Crohn’s disease and ulcerative colitis. Both genetic (Cho 2008) and functional studies (Xavier and Podolsky 2007) strongly support a role for STAT3 in IBD; indeed, STAT3 is a proposed therapeutic target in IBD (Lee et al. 2015; Nguyen et al. 2015). Our analysis also newly implicates FOXB1 in regulation of IBD genes. We compared the centrality of STAT3 and FOXB1 in the Th17 TRN (**Supplemental Fig. S20A**) to their centrality in the subnetwork limited to the 54 IBD genes in the Th17 TRN (**<u>Fig. 6B</u>**<u>, left panel</u>). In addition to degree centrality, we also examined betweenness centrality. For each TF, betweenness is the fraction of shortest paths connecting TFs to target genes in the network that contains the TF. Whereas degree is a local measure (TF’s direct effect on gene expression), betweenness is a more global measure of TF importance as it can also capture TFs that regulate a large number of genes through control of other TFs. Although STAT3 had the sixth-highest degree in the full Th17 TRN (**Supplemental Fig. S20A**), it becomes the highest-degree TF in the IBD subnetwork (**Fig. 6B**). Relative degree more than doubles for both STAT3 and FOXB1 in the IBD subnetwork, and betweenness centrality increased, too (Fig. 6B). The IBD genes regulated by STAT3 and FOXB1 include a number of Th17 genes: *Rorc, Il23r, Tnfsf15* (**<u>Fig. 6B</u>**<u>, right panel</u>).

**Figure 6.**
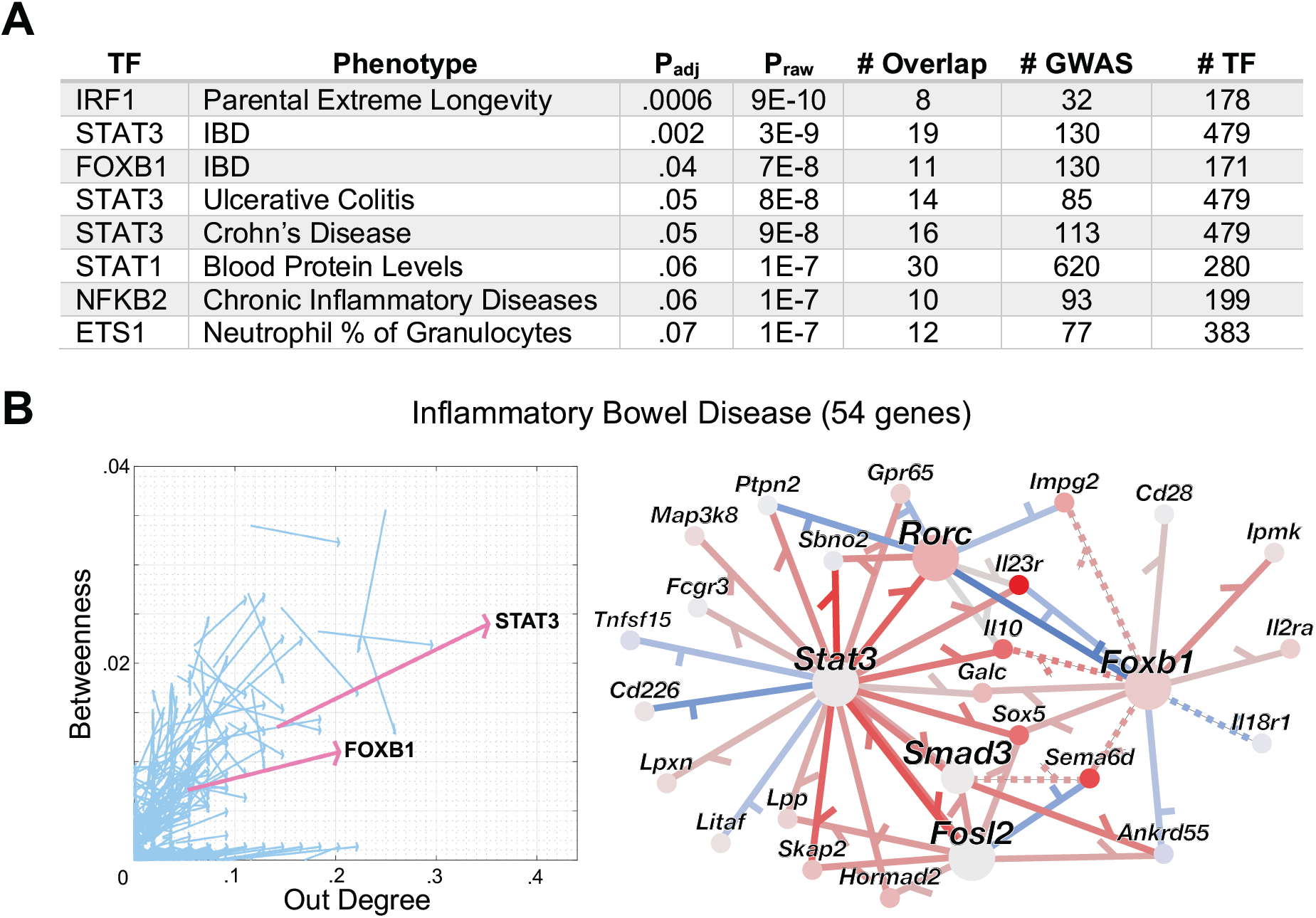
(A) TFs whose target genes are enriched in GWAS phenotype genes (FDR=10%). Phenotype abbreviations “IBD” = “Inflammatory Bowel Disease” and “Chronic inflammatory diseases” = “Chronic inflammatory diseases (ankylosing spondylitis, Crohn’s disease, psoriasis, primary sclerosing cholangitis, ulcerative colitis) (pleiotropy)”. # GWAS, # TF, and # Overlap correspond to the number of genes associated with the phenotype, regulated by the TF in the Th17 TRN (KO+ChIP+ATAC), and the overlap between those two sets, respectively. Further details are contained in **<u>Methods</u>**. **(B) STAT3 and FOXB1 are central regulators of IBD genes**. In the left panel, each arrow corresponds to a single TF. Arrow source is TF’s centrality (out degree, betweenness) in the full Th17 TRN and arrowhead is TF centrality for the IBD subnetwork (where target genes are limited to the 54 shared the Th17 TRN and IBD GWAS set). STAT3 and FOXB1 (pink arrows) both show significant increase in degree centrality for IBD genes (FDR=10%). The right panel features the subnetwork connecting STAT3 and FOXB1 to their target genes in the IBD set. Node color indicates log2(fold-change) in Th17 48h condition relative to other Th timepoints (red = increased, blue = decreased), while red / blue indicate positive / negative regulation. Solid edges have support in the ChIP+KO+ATAC prior, while dotted edges do not.

NFKB2 and ETS1 are also significantly associated with immune phenotypes (**Fig. 6A**). NFKB2’s targets are enriched in the phenotype “Chronic inflammatory diseases (ankylosing spondylitis, Crohn’s disease, psoriasis, primary sclerosing cholangitis, ulcerative colitis) (pleiotropy)” (**Supplemental Fig. S20B**). Mutations in *NFKB2* have been previously associated with common variable immunodeficiency (CVID) (Liu et al. 2014; Chen et al. 2013; Lindsley et al. 2014), a heterogeneous disorder in which 25% of patients suffer autoimmune disorders, including thrombocytopenia purpura, autoimmune hemolytic anemia, rheumatoid arthritis, and autoimmune enteropathy (which can be classified as Crohn’s disease) (Cunningham-Rundles 2008; Lopez-Herrera et al. 2012). Thus, NFKB2 is already genetically associated with pleiotropic autoimmune diseases, in the context of CVID. (We note that our set of GWAS phenotypes did not include CVID.) ETS1’s targets are associated with “Neutrophil percentage of granulocytes” (**Supplemental Fig. S20C**). ETS1 is known to repress the Th17 program (Moisan et al. 2007). *Ets1* expression decreases over the course of both Th0 and Th17 polarization, and, of the 48h Th polarization conditions, has highest expression in T_reg_. Mutations in *ETS1* have been associated with systemic lupus erythematosus (SLE) (Leng et al. 2011), an autoimmune disease in which the role of neutrophils has become increasingly appreciated (Smith and Kaplan 2015). Thus, a predicted role in neutrophil regulation could be consistent with the known role of ETS1 in SLE.

## Discussion

Th17 cells protect mucosa from bacteria and fungi but can also drive autoimmune and inflammatory disease (Khader et al. 2009; Littman and Rudensky 2010; Stadhouders et al. 2017). These diverse roles require coordination of thousands of genes. TF-specific knowledge of gene expression regulation is of great interest, providing a map for immuno-engineering Th17 behavior in disease. Researchers in academia and industry have used our first genome-scale Th17 transcriptional regulatory network (Ciofani et al. 2012) to develop hypotheses in the context of autoimmunity (Patel and Kuchroo 2015; Yang et al. 2014; Isono et al. 2014). Here, we provide an important update to our knowledge of Th17 transcriptional regulation, enabled by technical advances in genomic measurement and computational advances in TRN inference. KO data for 20 TFs and TF ChIP data for 9 TFs were central to the original Th17 TRN, providing excellent coverage of TF-gene targets for TFs in that set and a resource for the Th17 community. However, technical limitations and cost precluded application of these tools to the hundreds of TFs expressed over the course of Th17 differentiation, all of which could play important roles in Th17 gene expression regulation.

In combination with large-scale efforts to learn TF DNA-binding motifs (Weirauch et al. 2014; Jolma et al. 2013; Najafabadi et al. 2015; Badis et al. 2009), the advent of ATAC-seq represents an opportunity to overcome limitations of sequential TF ChIP experiments, expanding the number of TFs with chromatin binding profiles by over an order of magnitude (**Supplemental Table S1B**). In addition, although TF KO and ChIP data were pragmatically limited to 48h Th17 conditions, standard ATAC-seq protocols require two orders of magnitude fewer cells than TF ChIP. Here, we were able to obtain (indirect) TF binding profiles from multiple Th17 differentiation time points. Yet TF-binding profiles derived from motif analysis of ATAC-seq are noisy, containing false positives and false negatives. For example, in this study, only ~40% of mouse TFs had known motifs, so motif analysis of ATAC-seq data excluded more than half of potential TF regulators. Here, we provide a single, integrated method to infer regulatory roles for TFs genome-wide. At its core, gene expression is modeled as a function of TF activities, where prior information (e.g., from ATAC-seq) can be used to (1) improve TF activity estimates (and therefore target prediction) for some TFs and/or (2) favor TF-gene interactions that also have support from other sources (e.g., ATAC-seq, KO). We rigorously test the performance of our method in terms of precision-recall of gold-standard interactions from TF KO and ChIP and verify TRN model quality in terms of gene-expression prediction. Our methods have two very desirable features (1) they prune initial noisy prior networks (by over an order of magnitude in this study) while (2) also learning new TF-gene interactions for TFs with and without prior information.

Our final Th17 TRN is built integrating our best knowledge for model-building (combining knock-out of key TFs, ChIP-seq of multiple Th17-relevant TFs, and ATAC-seq with a rich gene expression dataset). While the original “core” Th17 network was limited to TFs with ChIP and KO data, our de novo Th17 core includes the original core (RORC, STAT3, BATF, IRF4, MAF) and dozens of additional TFs (**Fig. 5A**). We performed extensive gene-set enrichment to determine functional roles and potential phenotypes for novel core TFs. We detect TF-TF modules and make multivariate predictions for gene pathways regulated by multiple TFs. Although HIF1A was detected as an important TF regulating hypoxia-related genes in the original Th17 TRNs, here we strongly implicate several other factors (HIF3A, DPF1, SCRT1) in regulation of hypoxia genes. In addition, we uncover multivariate regulation of amino acid transport, interferon, and many other pathways (Fig. 5C, **S16-S19**). We also exhaustively test for the association of TF target genes with nearly 1000 GWAS phenotypes, uncovering known associations between STAT3 and IBD as well as several novel associations involving autoimmune and immune phenotypes (Fig. 6, **S20**). This work provides an important update to our knowledge of transcriptional regulation in Th17 cells and can be used to query key regulators of pathways and disease genes in Th17.

Of perhaps greater importance, the TRN experimental design and computational methods proposed are general, designed for regimes where prior knowledge of transcriptional regulators and/or sample material is scarce (e.g., cells directly from humans and animal models). Given the rigorous testing and case study presented here, we have high expectations for their successful application in other systems. Indeed, de novo Th17 cores and TF-TF modules are remarkably similar between TRNs derived from all prior information (KO+ChIP+ATAC) and ATAC-seq data only. In a companion study, we apply our methods to a physiological setting, constructing and experimentally validating TRNs for innate lymphoid cells of the intestine (Miraldi et al.). Our methods are general and widely applicable. Prior information need not be limited to motif analysis of ATAC-seq data. TF binding patterns can also be inferred from TF ChIP, histone-mark ChIP, FAIRE-seq, Dnase1-seq. In addition, prior information can be derived from orthogonal sources: systems genetics and perturbation screens as well as curated from the literature.

This work also highlights avenues for future improvement of TRN inference methods. We tested two methods for TF activity estimation: (1) TF mRNA levels and (2) based on prior knowledge of TF target gene expression. Although prior-based TFA improved TRN inference in B. subtilis and yeast (Arrieta-Ortiz et al. 2015; Tchourine et al. 2018), neither method consistently outperformed the other in this study. As a result, final TRNs were built using both TFA methods. There are multiple dimensions along which TFA estimation could be improved. The simplicity of the linear framework proposed for prior-based estimation has limitations in the context of complex mammalian transcriptional regulation, and a more sophisticated mathematical model for TFA estimation could be of value. TFA estimation would also improve from better prediction of TF binding events. Here, we limited our approach to a simple TF motif analysis of accessible chromatin, yet several more sophisticated methods exist and merit testing (Pique-Regi et al. 2011; Sherwood et al. 2014; Chen et al. 2017; Lamparter et al. 2017). Another limitation of our method is the crude way in which putative TF binding events are mapped to gene loci. In our analysis, 3D distance between potential regulatory regions and gene loci is approximated by linear distance, a short-coming that chromatin capture data (e.g., Hi-C (Lieberman-Aiden et al. 2009) and other 3D-chromatin techniques (Zhang et al. 2012; Beagrie et al. 2017) would mitigate. Thus, the Th17 genomics dataset (Ciofani et al. 2012), augmented by our new ATAC-seq and RNA-seq experiments, provides a fertile testing ground for the development of future TRN inference methods and innovation.

### Methods ATAC-seq

CD4+ T Cells were sorted and polarized according to (Ciofani et al. 2012), and ATAC-seq samples were prepared as described previously (Buenrostro et al. 2013). Paired-end 50bp sequences were generated from samples on an Illumina HiSeq2500. Sequences were mapped to the murine genome (mm10) with bowtie2 (2.2.3), filtered based on mapping score (MAPQ > 30, Samtools (0.1.19)), and duplicates removed (Picard). The ATACseqQC package (Ou et al. 2018) was used to evaluate ATAC-seq fragment-length distributions and signal at TSS for each sample (**Supplemental Fig. S1**). For each sample individually, we ran Peakdeck (McCarthy and O’Callaghan 2014) (parameters –bin 75, -STEP 25, -back 10000, -npBack100000) and filtered peaks with a P_raw_<1E-4. To enable quantitative comparison of accessibility across samples, we generated a reference set of accessible regions, taking the union (Bedtools) of peaks detected in individual samples. The reference set of ATAC-seq peaks contained 63,049 potential regulatory loci, ranging from 75 base pairs (bp) to 3725 bp (median length 275 bp). Reads per reference peak were counted with HTSeq-count. ATAC-seq data was robustly normalized using DESeq2 (Love et al. 2014) for PCA and clustering (Fig. 1A, **S2**). The 33 ATAC-seq experiments are available from NCBI’s GEO Database under accession GSE113721.

### RNA-seq

#### Data Sources

The 18 new samples composing “late Th17” time points were generated as follows: Naïve CD4 T cells were primed on an anti-CD3/CD28 coated plate (without any additional cytokine) for 12-14hrs (overnight) and then polarized the cells with one of two cytokine cocktails: (1) Th17N: TGF-b (0.3ng/ml) + IL6 (20ng/ml) and (2) Th17P: IL6 (20ng/ml) + IL1b (20ng/ml) + IL23 (20ng/ml). Cells were harvested for RNA-seq at 60 and 108hrs post-TCR stimulation. Cells were lysed and snap-frozen in trizol then thawed for chloroform extraction, and 70% ethanol was mixed in a 1:1 ratio with the aqueous phase. Samples were then loaded onto a Qiagen RNEasy column according to manufacturer’s instructions. A Ribo-Zero Gold kit was used to deplete rRNA; libraries were then prepared using the Illumina TruSeq Stranded Total RNA library prep and sequenced on an Illumina HiSeq2500. The remaining 81 new CD4+T cell samples were (1) naïve or polarized and (2) processed as described in (Ciofani et al. 2012). The 99 RNA-seq experiments are available from NCBI’s GEO Database under accession number GSE113720. Publicly available RNA-seq data was downloaded from GEO: GSE40918 (156 samples), GSE70108 (4 samples), and GSE92992 (8 samples).

#### Batch Effects

The final gene expression matrix contained samples generated over an eight-year period, from several genomics cores, and with a combination of 36-bp and 50-bp single-end reads as well as 50-bp paired-end reads. There was no common set of control samples across batches. We first attempted to control for differences in sequencing, mapping only a single-end from paired-end sequencing samples with STAR aligner (Dobin et al. 2013). In addition, we controlled for differences in mappability between 36-bp and 50-bp reads by calculating effective gene lengths. Specifically, we calculated the number of possible 36-mers or 50-mers that HTSeq-count would map per gene (using HTSeq-count parameters: --stranded=no --mode=union and 36-mer and 50-mer mappability tracks (Derrien et al. 2012)). We then computed a factor (effective 36-mer gene length to effective 50-mer gene length) to convert 50-mer gene quantification by HTSeq-count to 36-mer gene quantification scale. From there, we used DESeq2 (Love et al. 2014) to normalize data. We used PCA to ensure that major trends did not correlate with (1) sequencing, (2) date of experiment, or (3) library size. Instead, they correlated well with the T Cell time course and polarization conditions (Fig. 1A, **S3**).

### Transcriptional regulatory network inference

#### Selection of Target Genes

We built gene expression models for <u>3578</u> target genes, composed of the union of (1) genes differentially expressed between Th17 and Th0 at 48h (FDR = 10%, log_2_|FC| > log_2_(1.5)) and (2) the 2100 genes in the original Th17 TRN (Ciofani et al. 2012). This list is available in **Supplemental Table S2**.

#### Selection of Potential Regulators

We initially generated a custom list of potential mouse protein TFs, combining (1) mouse and human TFs from TFClass (Wingender et al. 2014) and genes with the GO annotation “Transcription Factor Activity”. (Human TFs were mapped to mouse using the MGI database.) From our list of potential mouse protein TFs (2093 genes), we generated a final list of <u>869</u> potential TF regulators, limited to those TFs with differential gene expression in at least one pairwise comparison between Th17, Th0, Th1, Treg, or Th2 at 48h (FDR = 10%, log_2_|FC| > log_2_(1.5)). This initial list of 869 TFs was used for all analyses comparing mLASSO-StARS and BBSR-BIC (Fig. 2, 3B, **S4, S6**). However, given very recent efforts in TF annotation (Lambert et al. 2018), we generated a new list of mouse TFs for all subsequent TRN analyses. Specifically, *Lambert et al*. began with a list of candidate human TFs and manually curated whether available data supported a TF assignment, yielding both lists of likely TFs and likely non-TFs. We again used MGI to convert both lists to mouse. To gain mouse TFs without human orthologs, we integrated with mouse TFs from AnimalTFDB (Zhang et al. 2015), but removed any mouse TFs (70) that were unlikely to be TFs (Lambert et al. 2018). Our final mouse TF list contained 1577 TFs, <u>715</u> of which served as potential regulators (differentially expressed as described above). Both lists of candidate TFs are available in **Supplemental Table S3**. Note that **Supplemental Tables S1A** and **S1B** correspond to the initial and updated TF lists, respectively.

#### Generation of Prior Matrices

*ATAC-seq*: ATAC-seq peaks were detected (as described above) and associated with putative transcription-factor (TF) binding events and target genes to generate a “prior” network,) *p*∈ℝ^|genes|×|TFs|^, of TF-gene interactions. We used a compendium of human and mouse TF motifs. Human and/or mouse transcription factor binding motifs (PWMs) were downloaded from the CisBP motif collection version 1.02 (http://cisbp.ccbr.utoronto.ca) (Weirauch et al. 2014) and the ENCODE motif collection (http://compbio.mit.edu/encode-motifs) (Kheradpour and Kellis 2014). Transfac version 2014.2 motifs (Wingender 2008) referenced in the human CisBP collection were converted from transfac frequency matrix format to .meme format with the MEME Suite tool transfac2meme version 4.10.1. To add human ENCODE motifs to the collection, redundant entries for each TF were first removed from the ENCODE set if the PWM was highly correlated (R^2^ > 0.95) with any CisBP entry for that TF. Human TFs in the combined ENCODE and CisBP set were then mapped to mouse orthologs. We scanned peaks for individual motif occurrences with FIMO (Grant et al. 2011) (parameters --thresh .00001, --max-stored-scores 500000, and a first-order-Markov background model). We found inclusion of human TF orthologs from the ENCODE motif collection slightly increased precision-recall relative to mouse CisBP alone (data not shown). TF motif occurrences with raw p-value < 1E-5 were included in downstream analysis. Putative binding events were associated with a target gene, if the peak fell within +/-10kb of gene body. We tested several peak-gene association rules based on distance from gene body or TSS, and TRN inference was robust to that choice (data not shown). We generated two ATAC-seq priors: (1) A(Th17) for which only peaks from Th17 48h wildtype conditions were included and (2) A(Th) for which all Th samples were included.

*ChIP-seq*: TF ChIP-seq and control sequencing data were downloaded from GEO (GSE40918), mapped to the murine genome (mm10) with bowtie2 (2.2.3), filtered based on mapping score (MAPQ > 30, Samtools (0.1.19)), and duplicates removed (Picard). Peaks were called with macs14 version 1.4.2 (parameters: -m 10,30 -g 1865500000 --bw=200) and retained for raw p-value < 10^-10^. TFs were associated with a target gene, if the ChIP peak fell within +/-10kb of gene body. We tested several peak-gene association rules based on distance from gene body or TSS as well as a raw p-value < 10^-5^ cutoff; downstream methods were robust to these decisions (data not shown).

*ENCODE DHS*: TF-gene interactions, based on TF-footprinting analysis in ENCODE DNase1-hypersensitivity (DHS) in 25 mouse tissues (Stergachis et al. 2014) were downloaded from http://www.regulatorynetworks.org.

*TRRUST*: Signed, human TF-gene interactions were downloaded from the Transcriptional Regulatory Relationships Unraveled by Sentence-base Text-mining (TRRUST) database (Han et al. 2015), version 1, and mapped to mouse orthologs using the MGI database (one-to-many mappings included).

*ChIP/A(Th17)*: This prior uses TF ChIP data when available and TF motif analysis of 48h Th17 ATAC-seq data otherwise.

*ChIP/A(Th17)+KO+A(Th)*: This prior represents a combination of ChIP/A(Th17) prior, the 25 TF KO experiments in the G.S., and A(Th) prior. Each prior matrix was Frobenius-norm normalized and multiplied by the number of TFs, so that the predictions of each TF per prior had comparable weight. The interactions in the normalized priors were then summed, and the sign of prior interaction was determined from KO data, when available.

Given the degeneracy of TF motifs, some TFs had identical target gene sets within ATAC-seq and ENCODE DHS priors. Degenerate TFs were merged for TRN inference. For precision-recall analysis, resulting target edges for merged TFs were included for all degenerate TFs in the set.

#### Inference framework

We used the *Inferelator* algorithm for TRN inference (Bonneau et al. 2006). At steady state, gene expression is modeled as a sparse, multivariate linear combination of TF activities (TFAs):

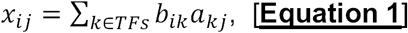

where x_ij_ corresponds to the expression level of gene i in condition j, a_jk_ is the activity of TF k in condition j, and b_ik_ is describes the effect of TF k on gene i. In the past, a TF’s mRNA expression level served as a proxy of protein TF activity. More recently (Arrieta-Ortiz et al. 2015), TFA has been estimated based on partial prior knowledge of a TF’s gene targets with the following equation:

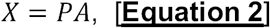

where *P* ∈ ℝ^|genes|×|TFs |^ is the prior matrix of known TF-gene interactions, *X* ∈ ℝ^|genes |×|samples|^ is the expression matrix for genes in the prior, and *A* ∈ ℝ^|TFs |×|samplesx^007^C;^ contains the unknown activity levels for TFs in the prior. There is no unique solution to Equation 2, but the least-squares solution has worked in practice (Arrieta-Ortiz et al. 2015; Tchourine et al. 2018). For this study, we tested both methods of TFA estimation: (1) TF mRNA levels and (2) based on prior knowledge of target genes (Equation 2). Note that the full set of expressed genes (24007) with edges in the prior were used when solving Equation 2. As described in results, we solved for the interaction terms {b_ik_} in Equation 1 using the current *Inferelator* engine (BBSR-BIC) as well as a new method (mLASSO-StARS, described in detail below).

#### Consideration of time-series samples

Our gene expression matrix includes a small fraction of time-series samples (15 out of 254), and so we tested whether linear differential equations, as proposed in the original *Inferelator* algorithm (Bonneau et al. 2006), would improve inference over the steady-state assumption (Equation 1) for our particular experimental design. For prior-based TFA (in contrast to TF mRNA), there is no obvious biological motivation for time-lag. However, given that TF mRNA is used to estimate the activities of TFs without prior information (~60% of TFs in our study), the steady-state assumption could also affect prior-based TFA. Using the time-lag parameter tau (30 minutes) from (Ciofani et al. 2012), we compared performance between TRNs built with and without time-lag. Precision-recall of the KO+ChIP and KO G.S.’s did not change between time-lagged and steady-state models (**Supplemental Fig. S21**). However, we do recommend linear modeling of differential gene expression for study designs involving more time-series data, and we provide workflows with this option in our codebase (https://github.com/emiraldi/infTRN_lassoStARS.git).

#### Model-building with LASSO and StARS

We used a modified Least Absolute Shrinkage and Selection Operator (mLASSO) to solve for a (sparse) set of TF-gene interaction terms {b_ik_} in Equation 1:

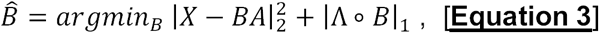

where X, and A are defined as above, B ∈ ℝ^|genes |×|TFs|^ is the matrix of inferred TF-gene interaction coefficients, Λ ∈ ℝ^|genes |×|TFs |^ is a matrix of nonnegative penalties, and ° represents a Hadamard (entry-wise matrix) product (Studham et al. 2014; Gustafsson et al. 2015). Representing the LASSO penalty as a Hadamard product involving a matrix of penalty terms, as opposed to a single penalty term, enabled us to incorporate prior information into the model-building procedure. Specifically, a smaller penalty Λ_*ik*_ is used if there is evidence for the TF-gene interaction in the prior matrix. Similar to the G-prior in the current *Inferelator* BBSR and older *Inferelator* modified elastic net (MEN) framework (Greenfield et al. 2013), this procedure encourages selection of interactions supported by the prior (e.g., containing ATAC-seq evidence), if there is also support in the gene expression data. For this study, the entries of the Λ matrices were limited to two values: the nonnegative valueλ, for TF-gene interactions without evidence in the prior, and bias*λ, where bias ∈ [0,1], for TF-gene interactions with support in the prior.

To choose the λ parameter, we tested the Stability Approach to Regularization Selection (StARS) (Liu et al. 2010). The method worked well for a closely related inference problem, functional gene networks (Zhang et al. 2016; Caballe Mestres et al. 2017), but, to our knowledge, this is the first application of StARS to model selection in the context of TRN inference and with incorporation of prior information as in Equation 3. StARS was designed to ensure that the inferred network of interactions includes the true set of network interactions with high probability. In contrast, another popular λ selection method, stability selection, seeks to limit false positive rate (Meinshausen and Bühlmann 2010), which, in a biological setting, might be overly conservative (Liu et al. 2010). Thus, we chose StARS over stability selection. The StARS authors describe their method as an “overselect” method, and this was essentially the behavior we desired to obtain gene models sufficiently large to explain more complex mammalian transcriptional patterns. This property was also key to the TF-TF module analysis describe below, as TF-TF modules were difficult to detect from smaller BBSR-BIC networks. In brief, StARS rests on the definition of edge instabilities. For a fixed value ofλ, instabilities are estimated via subsampling and can be interpreted as twice the Bernoulli variance of a subsampled edge or the fraction of times subsample edge predictions disagree. Authors provide advice about how to select a network with an acceptably (small) level of average edge instability and recommended a cutoff = .05.

We tested two ways of calculating average instability: for each gene model individually (“per-gene”) and for all gene models at a given λ (“network”). We reasoned that “per-gene” estimates might lead to better models for each gene, but the “network” average instability estimates would be more stable, requiring less search time for the λ corresponding to the desired instability cutoff (**Supplemental Fig. S22**). We visualized the distribution of subsampled TF-gene edge frequencies for per-gene and network average instability estimates at cutoffs = .05, .1 and .2 (**Supplemental Fig. S23**). Typically, the edges selected for the final network are those that are stably nonzero (i.e., present), with an instability less than or equal to the cutoff. For both per-gene and network average instability estimates, we noted that the average network size ranged from ~3.5 TFs/gene to ~9 TFs/gene for the default cutoff .05 to .2. Thus, conventional filtering of edges with StARS would lead to very sparse models, especially using the recommended .05 cutoff.

Importantly, given our application of StARS in the new setting of TRN inference and a modified LASSO objective function (matrix of penalties Λ) to integrate prior information, we used out-of-sample gene expression prediction and precision-recall of G.S. interactions to guide selection of an appropriate instability cutoff, rather than relying on the heuristic .05 cutoff. As we had done previously with application of StARS to ecological network inference (Kurtz et al. 2015), interactions were ranked for precision-recall according to nonzero subsamples per edge. However, this lead to a plateau at the beginning of the precision-recall curve (at high precision, low recall) for 50 subsamples (**Supplemental Fig. S24A**), and the plateau was not much improved by more time-consuming calculations involving 100 or 200 subsamples. To better and more efficiently prioritize high-confidence edges, we propose the following:

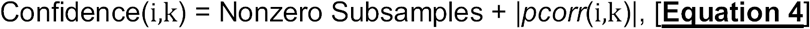

where i and k, again correspond to gene i and TF k, and *pcorr*(i,k) is the partial correlation between gene i and TF k, for a set model size. (Here, model size was set to 20 TFs/gene for partial correlation estimates; results were not sensitive to a range of sizes: 5-20 TFs/gene.) For TF-gene interactions with the same number of nonzero subsamples, the TF-gene interaction with high absolute partial correlation will be higher confidence. Equation 4 had the desired effect, improving precision-recall for both KO+ChIP (**Supplemental Fig. S24, S25**) and KO (**Supplemental Fig. S26**) gold standards (for both network-and gene-level average instabilities over a range of cutoffs: .05, .1 and .2). We also tested the same sets of conditions for all three out-of-sample prediction leave-out sets; representative results for the Early Th17 conditions are shown in **Supplemental Fig. S27**. Overall, precision-recall and out-of-sample prediction were robust to choice of network-versus per-gene average instability cutoff (**Supplemental Fig. S25, S26**). Out-of-sample prediction was consistently better using the instability cutoff .05 versus .1 and .2 (**Supplemental Fig. S27**). Thus, we chose the cutoff of .05, additionally noting reduced computation time that (.05 corresponds to a larger λ). Given that network-level average instability performed on-par with per-gene and was computationally less expensive, we used network-level estimates for all subsequence analyses.

Although we relied on the instability cutoff of .05 to select λand calculate edge confidences, we did not use the instability cutoff to select which edges to include in the final network, as is typically done using StARS. Instead, we recommend using quality metrics (precision, recall and out-of-sample prediction) to guide in selecting the final network size. Small differences in the cutoff (.05 to .2) led to large differences in network size (**Supplemental Fig. S23**), while network quality-metric curves (especially precision-recall (**Supplemental Fig. S25, S26**)), based only on ranking of TF-gene interactions (Equation 4), did not change much. However, applying the StARS cutoff to select the network size (represented by circles and x’s on curves in **Supplemental Fig. S25, S26**) is likely overly conservative. For example, applying cutoff = .05 yields a model size averaging 3 TFs/gene and corresponds to a median per-gene R^2^_pred_ value that is significantly lower than that of larger models (e.g., ~2-fold lower than median per-gene R^2^_pred_ at 15 TFs/gene, **Supplemental Fig. S27B**), suggesting that using cutoff = .05 to select model size would omit TF-gene interactions with predictive value from the model. Based on these analyses, we recommend using the above quality metrics to guide selection of model-size cutoff.

mLASSO-StARS was implemented in MATLAB R2016b, and code relies on the Glmnet for Matlab package (Qian et al. 2013) to solve Equation 3. To expedite instability calculations, we use average instability upper and lower bounds from bStARS (Müller et al. 2016). Using two subsamples to calculate upper and lower bounds and 50 subsamples for final instability estimates, the speed up from bStARS was negligible, while lower bounds estimated from five subsamples were tighter (**Supplemental Fig. S28**) and computation time was decreased > two-fold (from 75 to 35 minutes using a single core on a 2017 MacBook Pro with 2.9 GHz Intel Core i7 processor). Code is available at https://github.com/emiraldi/infTRN_lassoStARS.git and available in **Supplemental Materials**.

We tested several levels of prior reinforcement for both BBSR-BIC and mLASSO-StARS. For BBSR-BIC, G-prior weights of 1, 1.1, and 1.5 corresponded to no, moderate and strong levels of prior reinforcement, while, for mLASSO-StARS, bias = 1, .5, .25, (defined above) corresponded to no, moderate and strong reinforcement. These levels of prior reinforcement are similar between BBSR-BIC and mLASS-StARS in terms of resulting prior edges in the TRN (**Supplemental Fig. S5**).

#### Gene Expression Prediction

As described in **<u>Results</u>**, we generated three leave-out (test) datasets: “Early Th17” (all Th17 time points between 1-16 hours, 8 samples), “All Th0” (Th0 samples for all time points and perturbations, 53 samples), or “Late Th17” (18 Th17 samples from 60-108h post-TCR stimulation) (**Fig. 3A**). The samples corresponding to each leave-out set are listed in **Supplemental Table S4**. For each leave-out prediction challenge, the training set, as typical, was defined as all samples excluding test. For each training set, we performed **<u>both</u>** model selection and parameter estimation independently of the test set. Both BBSR-BIC and the mLASSO-StARS methods provide confidence estimates for predicted TF-gene interactions, and we built TRN models of various sizes as a function of edge confidence cutoffs for each of the training sets. For parameter estimation, training TFA matrices were mean-centered and variance-normalized according to the training-set means 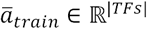 and standard deviations 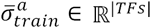 Target gene expression vectors were mean-centered according to the training-set mean 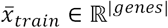. Then, for each confidence-level cutoff, we regressed the vector of normalized training gene expression data onto the reduced set of normalized training TFA estimates to arrive at a set of multivariate linear coefficients *B*_*train*_∈ℝ^|genes |×|TFs|^. Sum of Squared Error of prediction was calculated as follows:

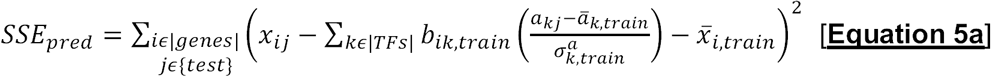

The “null” model *SSE* was calculated relative to the mean of training data:

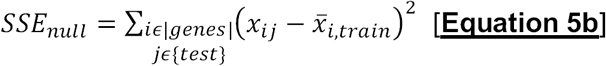

From *SSE*_*pred*_ and *SSE*_*null*_ we then calculated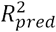, a normalized measure of predictive performance:

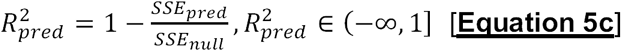

Intuitively, the 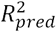compares prediction of the TRN model *SSE*_*pred*_(numerator) to the simplest model (based on mean of observed (training) data, *SSE*_*null,*_denominator). Any 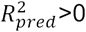indicates that the numerator model has some predictive benefit over the null model. For example, 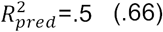means that the numerator model explains twice (thrice) more data variance than the intercept model.

For gene-expression prediction with prior-based TFA, the mRNA of target genes with edges in the prior contribute to TFA estimation (Equation 2), and then their gene expression patterns are predicted as a function of TFA. This circularity could lead to over-fitting and inflated 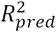values. To test this possibility, we constructed a prior-based TFA, A^-ij^, for each gene *i*, by solving:

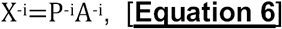

where matrices X^-i^, and P^-i^omit the rows corresponding to target gene i. Given the large gene-dimension of the prior (e.g., 17,138 genes for the ATAC (Th17) prior), we anticipated that A^-i^~ A. To test this assumption, we regenerated results for prior-based TFA (Th17 ATAC prior, bias = .5) according to [Equation 6]. There were 2803 target genes (out of 3578) with edges in the ATAC prior. For those TFs k that lost an edge to gene i when calculating A^-i^, we calculated the Pearson correlation between A^-i^ (k) and A(k) and plotted the cumulative distribution function (CDF) for the 108,850 resulting correlations (**Supplemental Fig. S29**). 99% of the TF activities were correlated with rho > .99, while a handful of correlations dipped below rho .95 (15 correlations (1.4%)). Thus, A^-i^ ~ A. We repeated leave-out gene expression prediction used gene-specific A^-i^, and, for all three expression prediction challenges, results were similar, even when limiting to the set of target genes with edges in the prior (**Supplemental Fig. S30**). This analysis confirms that our 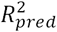estimates were robust to our decision to approximate A^-i^ ~ A.

#### Final TRNs and De Novo Th17 Core

Given the complementary performance of TF-mRNA-and prior-based TFA, we sought to combine the TRNs using both TFA methods (at moderate prior reinforcement). Given, their unique TF-specific AUPR profile (**Fig. 2C**), we combined methods by taking the maximum to preserve the individual strengths of each (Kittler et al. 1996; Castro et al. 2018) and compared to rank-sum (or average) combination (Marbach et al. 2012), which would emphasize commonalities but potentially mask individual strengths. As anticipated, max-rank combination had better precision-recall than rank-sum combination (**Supplemental Fig. S31**). For combination of edges with different sign, partial correlations were averaged to yield sign in the combined TRN. Given the orthogonal nature of the TRRUST prior (publicly available to those considering experimental designs for TRN inference), we also tested rank-combination of ATAC and TRRUST TRNs (combining a total of four TRNs, TF mRNA and prior-based TFA for each prior), but observed no benefit, in terms of improved precision-recall (**Supplemental Fig. S32**). More sophisticated methods for integrating TRNs would likely show benefit over the simple techniques tested here (Parisi et al. 2014; Castro et al. 2018), but are beyond the scope of this work.

De Novo Core Th17 TFs were limited to TFs specifically promoting Th17 gene expression patterns. TFs were included in the core if they met one of two criteria: (1) The TF promotes Th17 gene expression through activation (i.e., the TF’s positive edges are enriched in up-regulated Th17 genes at an FDR = 1%) or (2) repression of non-Th17 genes (i.e., TF’s negative edges are enriched in down-regulated Th17 genes at an FDR = 1%). The hypergeometric CDF and Benjamini-Hochberg procedure were used to estimate adjusted p-values.

#### Gold Standards

For the gold standards from our lab, we used recommended cutoffs of .75, .75, and 1.5 for KO, ChIP, and KO+ChIP networks, respectively (Ciofani et al. 2012). For the six additional TF KO experiments, we downloaded networks without filtering (Yosef et al. 2013). For both gold standards, gene symbols were mapped from mm9 to mm10, and only genes mapping to both genome builds were considered in precision-recall analysis.

*Random AUPR*. Random AUPR was calculated as the ratio of G.S. edges to target genes to the number of possible edges between target genes and TFs in the G.S.

### Network Visualization and Availability

Networks were visualized using a newly designed interactive interface, based on iPython and packages: igraph, numpy and scipy. The interface is still under development and will be detailed in a future publication; software is available at https://github.com/simonsfoundation/ jp_gene_viz. All 36 LASSO-StARS Th17 TRNs (from **Supplemental Fig. S10**), gold standards, and final, combined TRNs are available in a Jupyter-notebook binder: https://mybinder.org/v2/gh/simonsfoundation/Th17_TRN_Networks/ master. Both jp_gene_viz codebase and TRN notebooks are also included in **Supplemental Materials**.

### TF-TF Module Analysis

We calculated the number of shared target genes between each pair of TFs, analyzing positive and negative target edges separately. (Edges with |partial correlation| < .01 were excluded from analysis, as were TFs with fewer than 20 gene targets.) TFs vary greatly by number of target genes (**Supplemental Fig. S12, S13**), so we devised an overlap normalization scheme that controlled for the variable number of targets per TF. Specifically, we took inspiration from context likelihood of relatedness (CLR), a background-normalized mutual information score (Faith et al. 2007). We define the background-normalized overlap score *z*_*ij*_between TF *i* and TF *j* as:

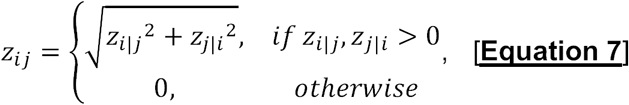

where *z*_*i|j*_is the z-score of the overlap between TF *i* and TF *j*, using the mean and standard deviation associated with the overlaps of TF *j* to calculate the z-score. Note that the score is only nonzero if the overlap is above average for both TFs. The normalized overlap score has good agreement with overlap significances estimated using the hypergeometric CDF.

We filtered the normalized overlap matrix so that it contained only TFs with at least one significant overlap (FDR = 10%, hypergeometric CDF). We then converted the similarity matrix of normalized overlaps to a distance matrix for hierarchical clustering using Ward distance. To arrive at a final number of clusters, we calculated the mean silhouette score for solutions over a range of total clusters and selected the solution that maximized mean silhouette score. For positive interactions (149 or 167 TFs), this analysis lead to 42 clusters, and 9 or 11 clusters for negative interactions (31 or 30 TFs), for final ChIP+KO+ATAC or ATAC-only TRNs, respectively.

To prioritize and rank TF-TF clusters, we developed a method based on combination of p-values generated from the empirical distribution of normalized overlap scores. For each cluster, we combined p-values for the pairwise normalized overlap scores between cluster TFs (e.g., for cluster of size *s*, there were *s*(*s*-1)/2 pairwise scores). We combined p-values with the weighted z-method (Whitlock 2005). P-values are converted via the inverse normal CDF to z-score space and combined with the following equation:

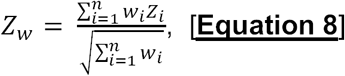

where *n* is the number of p-values to be combined and *w*∈ℝ^*n*^is a vector of weights. Z_*w*_is then converted to a p-value using the normal CDF. Here we set weights to 2/(*s* - 1). The resulting p-value score had the desired properties, ranking clusters on the basis of size and strength of overlap (**Supplemental Fig. S33A**) and was used to prioritize a limited set of 15 positive-edge TF-TF clusters for Fig. 5C, S16, S18. The significance of negative TF-TF clusters were orders of magnitude smaller than the top-15 positive TF-TF clusters (**Supplemental Fig. S33B**), so were not analyzed further. The lack of significant negative TF-TF clusters might have to do with the observed positive to negative edge bias in our method (~1.8-1.9:1).

### GWAS Analysis

The NHGRI-EBI GWAS Catalog v1.0.2 (MacArthur et al. 2016) was downloaded on August 4, 2018. SNPs were mapped to the nearest gene within +/-1Mbp using the catalog’s “MAPPED GENE(S)”. Phenotype-associated gene sets were then converted to mouse gene symbols and further filtered to gene sets containing five or more genes, resulting in 991 gene sets. For each TF in the Th17 TRN (KO+ChIP+ATAC prior) with five or more targets (605 TFs), overlap with the GWAS gene sets was calculated and significance estimated using the hypergeometric CDF. Benjamini-Hochberg correction was applied to control for multiple hypothesis testing. Network statistics were calculated in MATLAB R2016b, normalizing degree by total target genes and fraction of shortest paths (betweenness) by total number of paths between TFs and target genes (some of which were also TFs).

## Data Access

The newly generated RNA-seq and ATAC-seq data are available from NCBI’s GEO Database as SuperSeries GSE113723.

## Acknowledgements

We thank the Flatiron Institute Scientific Computing Core (I. Fisk) for enabling the computational aspects of this work and the New York University Langone Medical Center Genomics Core (A. Heguy and P. Zappile) for help with sequencing. We are thankful to T. Aijo, M. Weirauch and R. Taylor for advice on the manuscript and to G. Atluri for helpful discussions of the TF-TF module analysis. This work was supported by the Cincinnati Children’s Research Foundation (Trustee Award Grant to E.R.M.), the Simons Foundation (E.R.M., A.W., N.D., N.C., R.B.), U.S. National Institute of Health (5T32AI100853 to M.P.; R01-DK103358-01 to R.B. and D.L.; and R01-GM112192-01 to R.B., T32 CA009161 (Levy) to J.A.H.), Crohn’s and Colitis Foundation of America (fellowship to M.C.), Damon Runyon Cancer Research Foundation (Dale and Betty Frey Fellowship to J.A.H.), the Laura and Isaac Perlmutter Cancer Center (P30CA016087 to A.H.).

## References

Äijö T, Bonneau R. 2016. Biophysically Motivated Regulatory Network Inference: Progress and Prospects. Hum Hered 81: 62–77.

Arrieta-Ortiz ML, Hafemeister C, Bate AR, Chu T, Greenfield A, Shuster B, Barry SN, Gallitto M, Liu B, Kacmarczyk T, et al 2015. An experimentally supported model of the Bacillus subtilis global transcriptional regulatory network. Mol Syst Biol 11: 839.

Badis G, Berger MF, Philippakis AA, Talukder S, Gehrke AR, Jaeger SA, Chan ET, Metzler G, Vedenko A, Chen X. 2009. Diversity and complexity in DNA recognition by transcription factors. Science (80-) 324: 1720–1723.

Barski A, Cuddapah S, Cui K, Roh T-Y, Schones DE, Wang Z, Wei G, Chepelev I, Zhao K. 2007. High-Resolution Profiling of Histone Methylations in the Human Genome. Cell 129: 823–837.

Beagrie RA, Scialdone A, Schueler M, Kraemer DCA, Chotalia M, Xie SQ, Barbieri M, de Santiago I, Lavitas L-M, Branco MR, et al 2017. Complex multi-enhancer contacts captured by genome architecture mapping. Nature 543: 519.

Blatti C, Kazemian M, Wolfe S, Brodsky M, Sinha S. 2015. Integrating motif, DNA accessibility and gene expression data to build regulatory maps in an organism. Nucleic Acids Res 43: 3998–4012.

Bonneau R, Facciotti MT, Reiss DJ, Schmid AK, Pan M, Kaur A, Thorsson V, Shannon P, Johnson MH, Bare JC, et al 2007. A Predictive Model for Transcriptional Control of Physiology in a Free Living Cell. Cell 131: 1354–1365.

Bonneau R, Reiss DJ, Shannon P, Facciotti M, Hood L, Baliga NS, Thorsson V. 2006. The Inferelator: an algorithm for learning parsimonious regulatory networks from systemsbiology data sets de novo. Genome Biol 7: 1.

Boyle AP, Davis S, Shulha HP, Meltzer P, Margulies EH, Weng Z, Furey TS, Crawford GE. 2008. High-resolution mapping and characterization of open chromatin across the genome. Cell 132: 311–322.

Buenrostro JD, Giresi PG, Zaba LC, Chang HY, Greenleaf WJ. 2013. Transposition of native chromatin for fast and sensitive epigenomic profiling of open chromatin, DNA-binding proteins and nucleosome position. Nat Methods 10: 1213–8.

Buenrostro JD, Wu B, Litzenburger UM, Ruff D, Gonzales ML, Snyder MP, Chang HY, Greenleaf WJ. 2015. Singlecell chromatin accessibility reveals principles of regulatory variation. Nature 523: 486–90.

Caballe Mestres A, Bochkina N, Mayer C. 2017. Selection of the Regularization Parameter in Graphical Models using Network Characteristics. J Comput Graph Stat 0–0.

Castro DM, Veaux N de, Miraldi ER, Bonneau R. 2018. Multi-study inference of regulatory networks for more accurate models of gene regulation. bioRxiv 279224.

Cerami EG, Gross BE, Demir E, Rodchenkov I, Babur Ö, Anwar N, Schultz N, Bader GD, Sander C. 2010. Pathway Commons, a web resource for biological pathway data. Nucleic Acids Res 39: D685–D690.

Chai LE, Loh SK, Low ST, Mohamad MS, Deris S, Zakaria Z. 2014. A review on the computational approaches for gene regulatory network construction. Comput Biol Med 48: 55–65.

Chen K, Coonrod EM, Kumánovics A, Franks ZF, Durtschi JD, Margraf RL, Wu W, Heikal NM, Augustine NH, Ridge PG, et al 2013. Germline Mutations in NFKB2 Implicate the Noncanonical NF-kB Pathway in the Pathogenesis of Common Variable Immunodeficiency. Am J Hum Genet 93: 812–824.

Chen X, Yu B, Carriero N, Silva C, Bonneau R. 2017. Mocap: Large-scale inference of transcription factor binding sites from chromatin accessibility. Nucleic Acids Res 45: 4315–4329.

Cho JH. 2008. The genetics and immunopathogenesis of inflammatory bowel disease. Nat Rev Immunol 8: 458–466.

Choi GB, Yim YS, Wong H, Kim S, Kim H, Kim S V, Hoeffer CA, Littman DR, Huh JR. 2016. The maternal interleukin-17a pathway in mice promotes autism-like phenotypes in offspring. Science (80-) 351: 933 LP–939.

Christie D, Zhu J. 2014. Transcriptional regulatory networks for CD4 T cell differentiation. In Transcriptional Control of Lineage Differentiation in Immune Cells, pp. 125–172, Springer.

Ciofani M, Madar A, Galan C, Sellars M, Mace K, Pauli F, Agarwal A, Huang W, Parkurst CN, Muratet M, et al 2012. A validated regulatory network for Th17 cell specification. Cell 151: 289–303.

Consortium GO. 2004. The Gene Ontology (GO) database and informatics resource. Nucleic Acids Res 32: D258–D261.

Cunningham-Rundles C. 2008. Autoimmune manifestations in common variable immunodeficiency. J Clin Immunol 28: 42–45.

Debnath M, Berk M. 2014. Th17 pathway-mediated immunopathogenesis of schizophrenia: Mechanisms and implications. Schizophr Bull 40: 1412–1421.

Derrien T, Estellé J, Sola SM, Knowles DG, Raineri E, Guigó R, Ribeca P. 2012. Fast computation and applications of genome mappability. PLoS One 7: e30377.

Dobin A, Davis CA, Schlesinger F, Drenkow J, Zaleski C, Jha S, Batut P, Chaisson M, Gingeras TR. 2013. STAR: Ultrafast universal RNA-seq aligner. Bioinformatics 29: 15–21.

Duren Z, Chen X, Jiang R, Wang Y, Wong WH. 2017. Modeling gene regulation from paired expression and chromatin accessibility data. Proc Natl Acad Sci 114: E4914–E4923.

Endo Y, Yokote K, Nakayama T. 2017. The obesity-related pathology and Th17 cells. Cell Mol Life Sci 74: 1231–1245.

Faith JJ, Hayete B, Thaden JT, Mogno I, Wierzbowski J, Cottarel G, Kasif S, Collins JJ, Gardner TS. 2007. Large-scale mapping and validation of Escherichia coli transcriptional regulation from a compendium of expression profiles. PLoS Biol 5: e8.

Fu Y, Jarboe LR, Dickerson JA. 2011. Reconstructing genome-wide regulatory network of E. coli using transcriptome data and predicted transcription factor activities. BMC Bioinformatics 12: 233.

Giresi PG, Kim J, McDaniell RM, Iyer VR, Lieb JD. 2007. FAIRE (Formaldehyde-Assisted Isolation of Regulatory Elements) isolates active regulatory elements from human chromatin. Genome Res 17: 877–885.

Grant CE, Bailey TL, Noble WS. 2011. FIMO: scanning for occurrences of a given motif. Bioinformatics 27: 1017–1018.

Greenfield A, Hafemeister C, Bonneau R. 2013. Robust data-driven incorporation of prior knowledge into the inference of dynamic regulatory networks. Bioinformatics 29: 1060–1067.

Gustafsson M, Gawel DR, Alfredsson L, Baranzini S, Bjorkander J, Blomgran R, Hellberg S, Eklund D, Ernerudh J, Kockum I, et al 2015. A validated gene regulatory network and GWAS identifies early regulators of T cell-associated diseases. Sci Transl Med 7: 1–10.

Han H, Shim H, Shin D, Shim JE, Ko Y, Shin J, Kim H, Cho A, Kim E, Lee T, et al 2015. TRRUST: a reference database of human transcriptional regulatory interactions. Sci Rep 5: 11432.

Harley ITW, Stankiewicz TE, Giles DA, Softic S, Flick LM, Cappelletti M, Sheridan R, Xanthakos SA, Steinbrecher KA, Sartor RB, et al 2014. IL-17 signaling accelerates the progression of nonalcoholic fatty liver disease in mice. Hepatology 59: 1830–1839.

Hecker M, Lambeck S, Toepfer S, van Someren E, Guthke R. 2009. Gene regulatory network inference: Data integration in dynamic models-A review. BioSystems 96: 86–103.

Heng TSP, Painter MW, Elpek K, Lukacs-Kornek V, Mauermann N, Turley SJ, Koller D, Kim FS, Wagers AJ, Asinovski N. 2008. The Immunological Genome Project: networks of gene expression in immune cells. Nat Immunol 9: 1091.

Isono F, Fujita-Sato S, Ito S. 2014. Inhibiting RORγt/Th17 axis for autoimmune disorders. Drug Discov Today 19: 1205–1211.

Jolma A, Kivioja T, Toivonen J, Cheng L, Wei G, Enge M, Taipale M, Vaquerizas JM, Yan J, Sillanpa MJ, et al 2010. Multiplexed massively parallel SELEX for characterization of human transcription factor binding specificities. 861–873.

Jolma A, Yan J, Whitington T, Toivonen J, Nitta KR, Rastas P, Morgunova E, Enge M, Taipale M, Wei G, et al 2013. DNA-binding specificities of human transcription factors. Cell 152: 327–339.

Kanehisa M, Goto S. 2000. KEGG: kyoto encyclopedia of genes and genomes. Nucleic Acids Res 28: 27–30.

Karwacz K, Miraldi ER, Pokrovskii M, Madi A, Yosef N, Wortman I, Chen X, Watters A, Carriero N, Awasthi A, et al 2017. Critical role of IRF1 and BATF in forming chromatin landscape during type 1 regulatory cell differentiation. Nature 18: 412–421.

Khader SA, Gaffen SL, Kolls JK. 2009. Th17 cells at the crossroads of innate and adaptive immunity against infectious diseases at the mucosa. Mucosal Immunol 2: 403–411.

Kheradpour P, Kellis M. 2014. Systematic discovery and characterization of regulatory motifs in ENCODE TF binding experiments. Nucleic Acids Res 42: 2976–2987.

Kittler J, Hater M, Duin RPW. 1996. Combining classifiers. Proc - Int Conf Pattern Recognit 2: 897–901.

Kurtz ZD, Müller CL, Miraldi ER, Littman DR, Blaser MJ, Bonneau RA. 2015. Sparse and Compositionally Robust Inference of Microbial Ecological Networks. PLoS Comput Biol 11: e1004226.

Lambert SA, Jolma A, Campitelli LF, Das PK, Yin Y, Albu M, Chen X, Taipale J, Hughes TR, Weirauch MT. 2018. The Human Transcription Factors. Cell 172: 650–665.

Lamparter D, Marbach D, Rueedi R, Bergmann S, Kutalik Z. 2017. Genome-Wide Association between Transcription Factor Expression and Chromatin Accessibility Reveals Regulators of Chromatin Accessibility. PLoS Comput Biol 13: 1–19.

Lee SY, Lee SH, Yang EJ, Kim EK, Kim JK, Shin DY, Cho M La. 2015. Metformin ameliorates inflammatory bowel disease by suppression of the stat3 signaling pathway and regulation of the between Th17/Treg Balance. PLoS One 10: 1–12.

Lee TI, Rinaldi NJ, Robert F, Odom DT, Bar-Joseph Z, Gerber GK, Hannett NM, Harbison CT, Thompson CM, Simon I, et al 2002. Transcriptional Regulatory Networks in <em>Saccharomyces cerevisiae</em> Science (80-) 298: 799 LP–804.

Leng RX, Pan HF, Chen GM, Feng CC, Fan YG, Ye DQ, Li XP. 2011. The dual nature of Ets-1: Focus to the pathogenesis of systemic lupus erythematosus. Autoimmun Rev 10: 439–443.

Li P, Spolski R, Liao W, Leonard WJ. 2014. Complex interactions of transcription factors in mediating cytokine biology in T cells. Immunol Rev 261: 141–156.

Liao JC, Boscolo R, Yang Y-L, Tran LM, Sabatti C, Roychowdhury VP. 2003. Network component analysis: reconstruction of regulatory signals in biological systems. Proc Natl Acad Sci U S A 100: 15522–7.

Liberzon A, Subramanian A, Pinchback R, Thorvaldsdóttir H, Tamayo P, Mesirov JP. 2011. Molecular signatures database (MSigDB) 3.0. Bioinformatics 27: 1739–1740.

Lieberman-Aiden E, Van Berkum NL, Williams L, Imakaev M, Ragoczy T, Telling A, Amit I, Lajoie BR, Sabo PJ, Dorschner MO. 2009. Comprehensive mapping of long-range interactions reveals folding principles of the human genome. Science (80-) 326: 289–293.

Lindsley AW, Qian Y, Valencia CA, Shah K, Zhang K, Assa’ad A. 2014. Combined Immune Deficiency in a Patient with a Novel NFKB2 Mutation. J Clin Immunol 34: 910–915.

Littman DR, Rudensky AY. 2010. Th17 and Regulatory T Cells in Mediating and Restraining Inflammation. Cell 140: 845–858.

Liu H, Roeder K, Wasserman L. 2010. Stability Approach to Regularization Selection (StARS) for High Dimensional Graphical Models. In Advances in Neural Information Processing Systems 23 (eds. J.D. Lafferty, C.K.I. Williams, J. Shawe-Taylor, R.S. Zemel, and A. Culotta), pp. 1432–1440, Curran Associates, Inc.

Liu Y, Hanson S, Gurugama P, Jones A, Clark B, Ibrahim MAA. 2014. Novel NFKB2 Mutation in Early-Onset CVID. J Clin Immunol 34: 686–690.

Lopez-Herrera G, Tampella G, Pan-Hammarström Q, Herholz P, Trujillo-Vargas CM, Phadwal K, Simon AK, Moutschen M, Etzioni A, Mory A, et al 2012. Deleterious Mutations in LRBA Are Associated with a Syndrome of Immune Deficiency and Autoimmunity. Am J Hum Genet 90: 986–1001.

Love MI, Huber W, Anders S. 2014. Moderated estimation of fold change and dispersion for RNA-seq data with DESeq2. Genome Biol 15: 550.

MacArthur J, Bowler E, Cerezo M, Gil L, Hall P, Hastings E, Junkins H, McMahon A, Milano A, Morales J. 2016. The new NHGRI-EBI Catalog of published genome-wide association studies (GWAS Catalog). Nucleic Acids Res 45: D896–D901.

Madar A, Greenfield A, Vanden-Eijnden E, Bonneau R. 2010. DREAM3: Network inference using dynamic context likelihood of relatedness and the inferelator. PLoS One 5.

Marbach D, Costello JC, Küffner R, Vega NM, Prill RJ, Camacho DM, Allison KR, Aderhold A, Bonneau R, Chen Y. 2012. Wisdom of crowds for robust gene network inference. Nat Methods 9: 796.

McCarthy MT, O’Callaghan CA. 2014. PeaKDEck: a kernel density estimator-based peak calling program for DNaseI-seq data. Bioinformatics 30: 1302–1304.

Meinshausen N, Bühlmann P. 2010. Stability selection. J R Stat Soc Ser B (Statistical Methodol 72: 417–473.

Miraldi ER, Pokrovskii M, Hall JA, Ochayon DE, Yi R, Chaimowitz NS, Seelamneni H, Carriero N, Watters A, Waggoner S, et al. Characterization of transcriptional regulatory networks that drive the identities and functions of intestinal innate lymphoid cells. (in preparation).

Moisan J, Grenningloh R, Bettelli E, Oukka M, Ho I-C. 2007. Ets-1 is a negative regulator of Th17 differentiation. J Exp Med 204: 2825–2835.

Müller CL, Bonneau R, Kurtz Z. 2016. Generalized Stability Approach for Regularized Graphical Models.

Najafabadi HS, Mnaimneh S, Schmitges FW, Garton M, Lam KN, Yang A, Albu M, Weirauch MT, Radovani E, Kim PM. 2015. C2H2 zinc finger proteins greatly expand the human regulatory lexicon. Nat Biotechnol 33: 555.

Neph S, Stergachis AB, Reynolds A, Sandstrom R, Borenstein E, Stamatoyannopoulos JA. 2012. Circuitry and Dynamics of Human Transcription Factor Regulatory Networks. Cell 150: 1274–1286.

Nguyen PM, Putoczki TL, Ernst M. 2015. STAT3-Activating Cytokines: A Therapeutic Opportunity for Inflammatory Bowel Disease? J Interf Cytokine Res 35: 340–350.

Ou J, Liu H, Yu J, Kelliher MA, Castilla LH, Lawson ND, Zhu LJ. 2018. ATACseqQC: a Bioconductor package for post-alignment quality assessment of ATAC-seq data. BMC Genomics 19: 169.

Ouyang Z, Zhou Q, Wong WH. 2009. ChIP-Seq of transcription factors predicts absolute and differential gene expression in embryonic stem cells. Proc Natl Acad Sci 106: 21521–21526.

Parisi F, Strino F, Nadler B, Kluger Y. 2014. Ranking and combining multiple predictors without labeled data. Proc Natl Acad Sci U S A 111: 1253–8.

Patel DD, Kuchroo VK. 2015. Th17 Cell Pathway in Human Immunity: Lessons from Genetics and Therapeutic Interventions. Immunity 43: 1040–1051.

Pico AR, Kelder T, Van Iersel MP, Hanspers K, Conklin BR, Evelo C. 2008. WikiPathways: pathway editing for the people. PLoS Biol 6: e184.

Pique-Regi R, Degner JF, Pai AA, Gaffney DJ, Gilad Y, Pritchard JK. 2011. Accurate inference of transcription factor binding from DNA sequence and chromatin accessibility data. Genome Res 21: 447–455.

Qian J, Hastie T, Friedman J, Tibshirani R, Simon N. 2013. Glmnet for Matlab.

Qin J, Hu Y, Xu F, Yalamanchili HK, Wang J. 2014. Inferring gene regulatory networks by integrating ChIP-seq/chip and transcriptome data via LASSO-type regularization methods. Methods 67: 294–303.

Ramirez RN, El-Ali NC, Mager MA, Wyman D, Conesa A, Mortazavi A. 2017. Dynamic Gene Regulatory Networks of Human Myeloid Differentiation. Cell Syst 4: 416–429.e3.

Ren B, Robert F, Wyrick JJ, Aparicio O, Jennings EG, Simon I, Zeitlinger J, Schreiber J, Hannett N, Kanin E, et al 2000. Genome-Wide Location and Function of DNA Binding Proteins. Science (80-) 290: 2306 LP–2309.

Rendeiro AF, Schmidl C, Strefford JC, Walewska R, Davis Z, Farlik M, Oscier D, Bock C. 2016. Chromatin accessibility maps of chronic lymphocytic leukaemia identify subtype-specific epigenome signatures and transcription regulatory networks. Nat Commun 7.

Robertson G, Hirst M, Bainbridge M, Bilenky M, Zhao Y, Zeng T, Euskirchen G, Bernier B, Varhol R, Delaney A, et al 2007. Genome-wide profiles of STAT1 DNA association using chromatin immunoprecipitation and massively parallel sequencing. Nat Methods 4: 651.

Sherwood RI, Hashimoto T, O’Donnell CW, Lewis S, Barkal AA, Van Hoff JP, Karun V, Jaakkola T, Gifford DK. 2014. Discovery of directional and nondirectional pioneer transcription factors by modeling DNase profile magnitude and shape. Nat Biotechnol 32: 171–178.

Siahpirani AF, Roy S. 2016. A prior-based integrative framework for functional transcriptional regulatory network inference. Nucleic Acids Res 45: gkw963.

Smith CK, Kaplan MJ. 2015. The role of neutrophils in the pathogenesis of systemic lupus erythematosus. Curr Opin Rheumatol 27: 448–453.

Stadhouders R, Lubberts E, Hendriks RW. 2017. A cellular and molecular view of T helper 17 cell plasticity in autoimmunity. J Autoimmun.

Stergachis AB, Neph S, Sandstrom R, Haugen E, Reynolds AP, Zhang M, Byron R, Canfield T, Stelhing-Sun S, Lee K, et al 2014. Conservation of trans-acting circuitry during mammalian regulatory evolution. Nature 515: 365–370.

Studham ME, Tjärnberg A, Nordling TEM, Nelander S, Sonnhammer ELL. 2014. Functional association networks as priors for gene regulatory network inference. Bioinformatics 30: 130–138.

Tchourine K, Vogel C, Bonneau R. 2018. Condition-Specific Modeling of Biophysical Parameters Advances Inference of Regulatory Networks. Cell Rep 23: 376–388.

Weirauch MT, Yang A, Albu M, Cote AG, Montenegro-Montero A, Drewe P, Najafabadi HS, Lambert SA, Mann I, Cook K, et al 2014. Determination and Inference of Eukaryotic Transcription Factor Sequence Specificity. Cell 158: 1431–1443.

Whitlock MC. 2005. Combining probability from independent tests: the weighted Z-method is superior to Fisher’s approach. J Evol Biol 18: 1368–73.

Wilkins O, Hafemeister C, Plessis A, Holloway-Phillips M-M, Pham GM, Nicotra AB, Gregorio GB, Jagadish SVK, Septiningsih EM, Bonneau R, et al 2016. EGRINs (Environmental Gene Regulatory Influence Networks) in Rice That Function in the Response to Water Deficit, High Temperature, and Agricultural Environments. Plant Cell 28: 2365–2384.

Wingender E. 2008. The TRANSFAC project as an example of framework technology that supports the analysis of genomic regulation. Brief Bioinform 9: 326–332.

Wingender E, Schoeps T, Haubrock M, Dönitz J. 2014. TFClass: a classification of human transcription factors and their rodent orthologs. Nucleic Acids Res 43: D97–D102.

Xavier RJ, Podolsky DK. 2007. Unravelling the pathogenesis of inflammatory bowel disease. Nature 448: 427–434.

Xi H, Shulha HP, Lin JM, Vales TR, Fu Y, Bodine DM, McKay RDG, Chenoweth JG, Tesar PJ, Furey TS. 2007. Identification and characterization of cell type–specific and ubiquitous chromatin regulatory structures in the human genome. PLoS Genet 3: e136.

Yang J, Sundrud MS, Skepner J, Yamagata T. 2014. Targeting Th17 cells in autoimmune diseases. Trends Pharmacol Sci 35: 493–500.

Yosef N, Shalek AK, Gaublomme JT, Jin H, Lee Y, Awasthi A, Wu C, Karwacz K, Xiao S, Jorgolli M, et al 2013. Dynamic regulatory network controlling TH17 cell differentiation. Nature 496: 461–8.

Zhang HM, Liu T, Liu CJ, Song S, Zhang X, Liu W, Jia H, Xue Y, Guo AY. 2015. AnimalTFDB 2.0: A resource for expression, prediction and functional study of animal transcription factors. Nucleic Acids Res 43: D76–D81.

Zhang J, Poh HM, Peh SQ, Sia YY, Li G, Mulawadi FH, Goh Y, Fullwood MJ, Sung W-K, Ruan X. 2012. ChIA-PET analysis of transcriptional chromatin interactions. Methods 58: 289–299.

Zhang XF, Ou-Yang L, Zhao XM, Yan H. 2016. Differential network analysis from cross-platform gene expression data. Sci Rep 6: 1–13.

